# COVID-19 research in Wikipedia

**DOI:** 10.1101/2020.05.10.087643

**Authors:** Giovanni Colavizza

## Abstract

Wikipedia is one of the main sources of free knowledge on the Web. During the first few months of the pandemic, over 5,200 new Wikipedia pages on COVID-19 have been created and have accumulated over 400M pageviews by mid June 2020.^1^ At the same time, an unprecedented amount of scientific articles on COVID-19 and the ongoing pandemic have been published online. Wikipedia’s contents are based on reliable sources such as scientific literature. Given its public function, it is crucial for Wikipedia to rely on representative and reliable scientific results, especially so in a time of crisis. We assess the coverage of COVID-19-related research in Wikipedia via citations to a corpus of over 160,000 articles. We find that Wikipedia editors are integrating new research at a fast pace, and have cited close to 2% of the COVID-19 literature under consideration. While doing so, they are able to provide a representative coverage of COVID-19-related research. We show that all the main topics discussed in this literature are proportionally represented from Wikipedia, after accounting for article-level effects. We further use regression analyses to model citations from Wikipedia and show that Wikipedia editors on average rely on literature which is highly cited, widely shared on social media, and has been peer-reviewed.

## 1 Introduction

Alongside the primary health crisis, the COVID-19 pandemic has been recognized as an information crisis, or an “infodemic” [66, 12, 23]. Widespread misinformation [57] and low levels of health literacy [44] are two of the main issues. In an effort to deal with them, the World Health Organization maintains a list of relevant research updated daily [69], as well as a portal to provide information to the public [2]; similarly does the European Commission [3], and many other countries and organizations. The need to convey accurate, reliable and understandable medical information online has never been so pressing.

Wikipedia plays a fundamental role as a public source of information on the Web, striving to provide “neutral” and unbiased contents [38]. Wikipedia is particularly important as go-point to access trusted medical information [57, 55]. Fortunately, Wikipedia biomedical articles have been repeatedly found to be highly visible and of high quality [5, 35]. Wikipedia’s verifiability policy mandates that readers can check the sources of information contained in Wikipedia, and that reliable sources should be secondary and published.^2^ These guidelines are particularly strict with respect to biomedical contents, where the preferred sources are, in order: systematic reviews, reviews, books and other scientific literature.^3^

The COVID-19 pandemic has put Wikipedia under stress with a large amount of new, often non-peer-reviewed research being published in parallel to a surge in interest for information related to the pandemic [18]. The response of Wikipedia’s editor community has been fast: since March 17 2020, all COVID-19-related Wikipedia pages have been put under indefinite sanctions entailing restricted edit access, to allow for a better vetting of their contents.^4^ In parallel, a WikiProject COVID-19 has been established and a content creation campaign is ongoing [18, 24].^5^ While this effort is commendable, it also raises questions on the capacity of editors to find, select and integrate scientific information on COVID-19 at such a rapid pace, while keeping quality high. As an illustration of the speed at which events are happening, in Figure 1 we show the average time in number of months from publication to a first citation from Wikipedia for a large set of COVID-19-related articles (see Section 3). In 2020, this time has gone to zero: articles on COVID-19 are frequently cited in Wikipedia immediately after or even before their official publication date, based on early access versions of articles.

**Figure 1:**
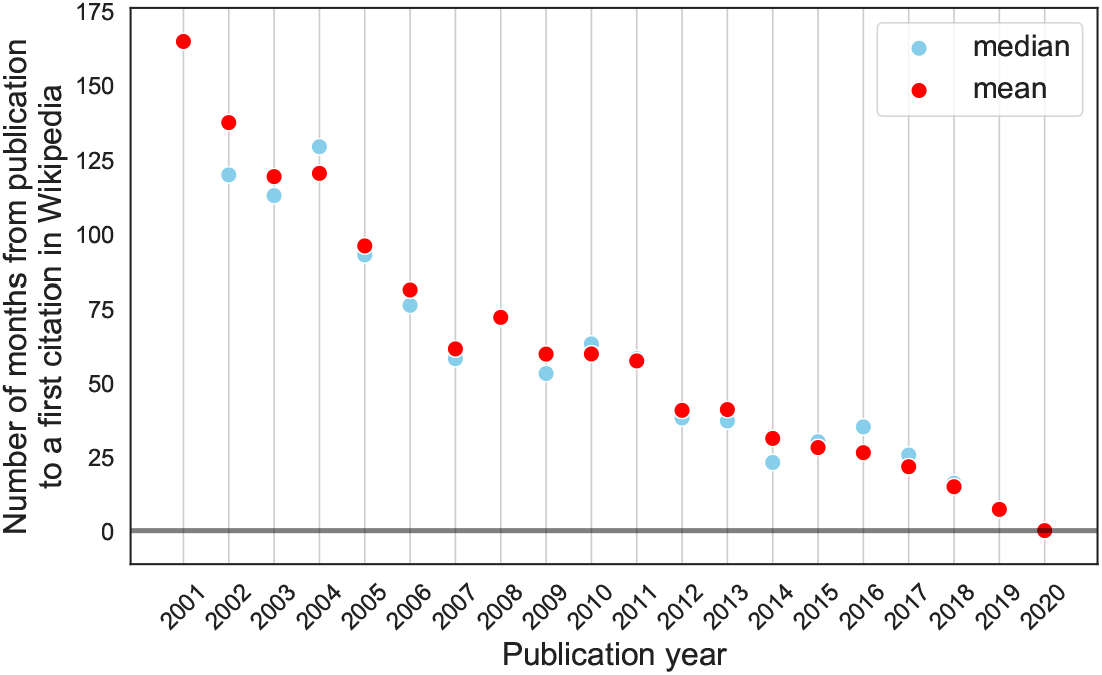
Number of months elapsed from publication to the first Wikipedia citation (mean and median binned by year) of COVID-19-related research. In 2020, the average number of months from (official) publication to the first citation from Wikipedia has gone to zero, likely due to the effect of early releases by some journals. Since this figure shows censored data, it should only be taken as illustrative of the fact that Wikipedia editors are citing very recent or even unpublished research.

In this work, we pose the following general question: *Is Wikipedia relying on a representative and reliable sample of COVID-19-related research?* We break this question down into the following two research questions:

1. RQ1: Is the literature cited from Wikipedia representative of the broader topics discussed in COVID-19-related research?
2. RQ2: Is Wikipedia citing COVID-19-related research during the pandemic following the same inclusion criteria adopted before and in general?

We approach the first question by clustering COVID-19-related publications using text and citation data, and comparing Wikipedia’s coverage of different clusters before and during the pandemic. The second question is instead approached using regression analysis. In particular, we model whether an article is cited from Wikipedia or not, and how many citations it receives from Wikipedia. We then again compare results for articles cited before and during the pandemic.

Our main finding is that Wikipedia contents rely on representative and high-impact COVID-19-related research. (RQ1) During the past few months, Wikipedia editors have successfully integrated COVID-19 and coronavirus research, keeping apace with the rapid growth of related literature by including a representative sample of each of the topics it contains. (RQ2) The inclusion criteria used by Wikipedia editors to integrate COVID-19-related research during the pandemic are consistent with those from before, and appear reasonable in terms of source reliability. Specifically, editors prefer articles from specialized journals or mega journals over pre-prints, and focus on highly cited and/or highly socially visible literature. Altmetrics such as Twitter shares, mentions in news and blogs, the number of Mendeley readers are complementing citation counts from the scientific literature as an indicator of impact positively correlated with citations from Wikipedia. After controlling for these article-level impact indicators, and for publication venue, time and size-effects, there is no indication that the topic of research matters with respect to receiving citations from Wikipedia. This indicates that Wikipedia is currently not over nor under-relying on any specific COVID-19-related scientific topic.

## 2 Related work

Wikipedia articles are created, improved and maintained by the efforts of the community of volunteer editors [48, 11], and they are used in a variety of ways by a wide user base [54, 32, 46]. The information Wikipedia contains is generally considered to be of high-quality and up-to-date [48, 25, 19, 30, 47, 5, 55], notwithstanding margins for improvement and the need for constant knowledge maintenance [11, 33, 17].

Following Wikipedia’s editorial guidelines, the community of editors creates contents often relying on scientific and scholarly literature [42, 20, 6], and therefore Wikipedia can be considered a mainstream gateway to scientific information [31, 21, 33, 52, 36, 46]. Unfortunately, few studies have considered the *representativeness and reliability* of Wikipedia’s scientific sources. The evidence on what scientific and scholarly literature is cited in Wikipedia is slim. Early studies point to a relative low overall coverage, indicating that between 1% and 5% of all published journal articles are cited in Wikipedia [49, 53, 68]. Previous studies have shown that the subset of scientific literature cited from Wikipedia is more likely on average to be published on popular, high-impact-factor journals, and to be available in open access [41, 59, 6].

Wikipedia is particularly relevant as a means to access medical information online [31, 21, 55, 57]. Wikipedia medical contents are of very high quality on average [5] and are primarily written by a core group of medical professionals part of the nonprofit Wikipedia Medicine [52]. Articles part of the WikiProject Medicine “are longer, possess a greater density of external links, and are visited more often than other articles on Wikipedia” [35]. Perhaps not surprisingly, the fields of research that receive most citations from Wikipedia are “Medicine (32.58%)” and “Biochemistry, Genetics and Molecular Biology (31.5%)” [6]; Wikipedia medical pages also contain more citations to scientific literature than the average Wikipedia page [36]. Margins for improvement remain, as for example the readability of medical content in Wikipedia remains difficult for the non-expert [10]. Given Wikipedia’s medical contents high quality and high visibility, our work is concerned with understanding whether the Wikipedia editor community has been able to maintain the same standards for COVID-19-related research.

## 3 Data and Methods

### 3.1 COVID-19-related research

COVID-19-related research is not trivial to delimit [15]. Our approach is to consider two public and regularly-updated lists of publications:

- The Dimensions COVID-19 Publications list [1].
- The COVID-19 Open Research Dataset (CORD-19): a collection of COVID-19 and coronavirus related research, including publications from PubMed Central, Medline, arXiv, bioRxiv and medRxiv [65]. CORD-19 also includes publications from the World Health Organization COVID-19 Database [4].

Publications from these three lists are merged, and duplicates removed using publications identifiers, including DOI, PMID, PMCID, Dimensions ID. Publications without at least one identifier among these are discarded. As of July 1, 2020, the resulting list of publications contains 160,656 entries with a valid identifier, of which 72,795 have been released in 2020, as it can be seen from Figure 2. The research on coronaviruses, and therefore the accumulation of this corpus over time, has been clearly influenced by the SARS (2003+), MERS (2012+) and COVID-19 outbreaks. We use this list of publications to represent COVID-19 and coronavirus research in what follows. More details are given in the online repositories.

**Figure 2:**
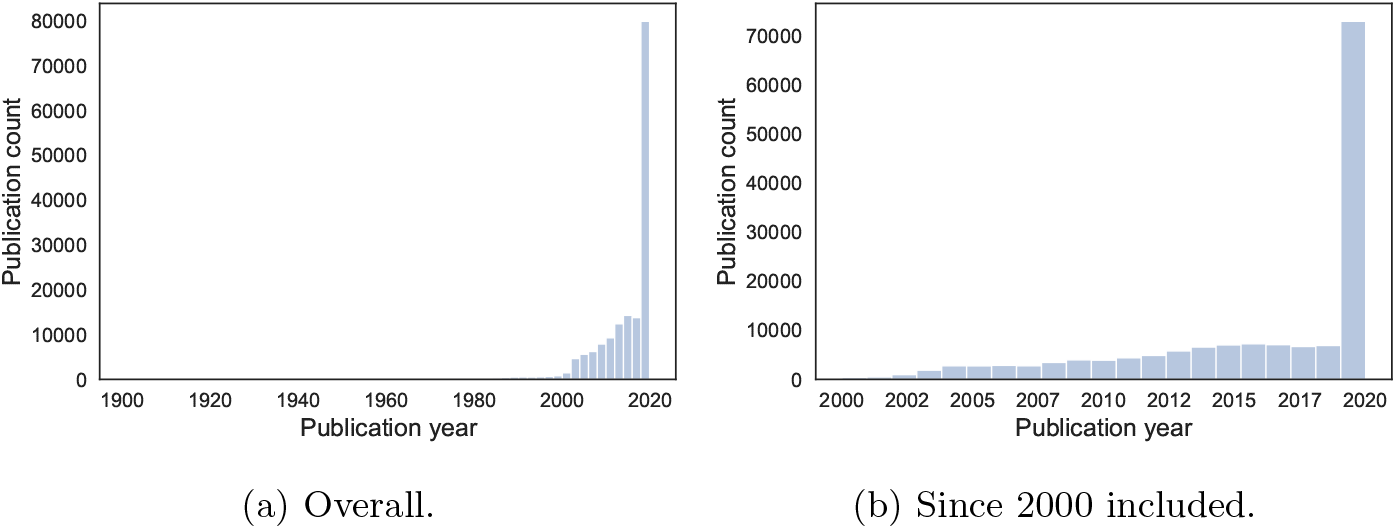
COVID-19-related literature over time, binned by publication year.

### 3.2 Auxiliary data sources

In order to study Wikipedia’s coverage of this list of COVID-19-related publications, we use data from Altmetric [51, 43]. Altmetric provides Wikipedia citation data relying on known identifiers.^6^ Despite this limitation, Altmetric data have been previously used to map Wikipedia’s use of scientific articles [68, 62, 6], especially since citations from Wikipedia are considered a possible measure of impact [56, 27]. Publications from the full list above are queried using the Altmetric API by DOI or PMID. In this way, 101,662 publications could be retrieved. After merging for duplicates by summing Altmetric indicators, we have a final set of 94,600 distinct COVID-19-related publications with an Altmetric entry.

Furthermore, we use data from Dimensions [22, 37] in order to get citation counts for COVID-19-related publications. The Dimensions API is also queried by DOI and PMID, resulting in 141,783 matches. All auxiliary data sources have been queried on July 1, 2020 too.

### 3.3 Methods

We approach our two research questions with the following methods:

1. RQ1: to assess whether the literature cited from Wikipedia is representative of the broader topics discussed in COVID-19-related research, we first cluster COVID-19 literature using text and citation data. Clusters of related literature allow us to identify broad distributions over topics within our COVID-19 corpus. We then assess to what extent the literature cited from Wikipedia follows the same distribution over topics of the entire corpus.
2. RQ2: to ascertain the inclusion criteria of Wikipedia editors, we use linear regression to model whether an article is cited from Wikipedia or not (logistic regression) and the number of Wikipedia citations it receives (linear regression).

In this section, we detail the experimental choices made for clustering analysis using publication text and citation data. Details on regression analyses are, instead, given in the corresponding section.

Text-based clustering of publications was performed in two ways: topic modelling and k-means relying on SPECTER embeddings. Both methods made use of the titles and abstracts of available publications, by concatenating them into a single string. We detected 152,247 articles in English, out of 160,656 total articles (−8409 over total). Of these, 33,301 have no abstract, thus we only used their title since results did not change significantly excluding articles without an abstract. Before performing topic modelling, we applied a pre-processing pipeline using scispaCy’s `en_core_sci_md` model [40] to convert each document into a bag-of-words representation, which includes the following steps: entity detection and inclusion in the bag-of-words for entities strictly longer than one token; lemmatisation; removal of isolated punctuation, stopwords and tokens composed of a single character; inclusion of frequent bigrams. SPECTER embeddings were instead retrieved from the API without any pre-processing.^7^

Topic modelling is a family of methods to learn statistical patterns of keywords frequently occurring together in the same documents. Formally, a topic is defined as a probability distribution over a vocabulary. Multiple topics can be learned from a corpus of documents and then used to cluster it [7]. While topic models are useful given that they require no annotated data, they also provide but a way to look at a certain corpus of documents. As such, they have been previously used for bibliometric analysis [67, 34]. We trained and compared topic models using Latent Dirichlet Allocation (LDA) [9], Correlated Topic Models (CTM) [8], Hierarchical Dirichlet Process (HDP) [58] and a range of topics between 5 and 50. We found similar results in terms of topic contents and in terms of their Wikipedia coverage (see Section 4) across models and over multiple runs, and a reasonable value of the number of topics to be between 15 and 25 from a topic coherence analysis [39]. Therefore, in what follows we discuss an LDA model with 15 topics.^8^ The top words for each topic of this model are given in the SI, while topic intensities over time are plotted as a heat map in Figure 8. SPECTER is a novel method to generate document-level embeddings of scientific documents based on a transformer language model and the network of citations [13]. SPECTER does not require citation information at inference time, and performs well without any further training on a variety of tasks. We embed every paper and cluster them using k-means with *k* = 20. The number of clusters was established using the elbow and the silhouette methods; different values of *k* could well be chosen, we again decided to pick the smallest reasonable value of *k*.

We then turned our attention to citation network clustering. We constructed a bibliographic coupling citation network [26] based all publications with references provided by Dimensions; these amount to 118,214. Edges were weighted using fractional counting [45], hence dividing the number of references in common between any two publications by the length of the union of their reference lists (thus, the max possible weight is 1.0). We only used the giant weakly connected component, which amounts to 114,829 nodes (−3385 over total) and 70,091,752 edges with a median weight of 0.0217. We clustered the citation network using the Leiden algorithm [64] with a resolution parameter of 0.05 and the Constant Potts Model (CPM) quality function [63]. With this configuration, we found that the largest 43 clusters account for half the nodes in the network, and the largest cluster is composed of 15,749 nodes.

These three methods differ in which data they use and how, and thus provide for complementary results. While topic models focus on word co-occurrences and are easier to interpret, bibliographic coupling networks rely on the explicit citation links among publications. Finally, SPECTER combines both kinds of data and modern deep learning techniques.

## 4 Results

Intense editorial work was carried out over the early weeks of 2020 in order to include scientific information on COVID-19 and coronaviruses into Wikipedia [24]. From Figure 3a, we can appreciate the surge in new citations added from Wikipedia to COVID-19 research. Importantly, these citations were not only added to cope with the growing amount of new literature, but also to fill gaps by including literature published before 2020, as shown in Figure 3b. The total fraction of COVID-19-related articles that are cited at least once from Wikipedia over the total is 1.9%. Yet, this number is uneven over languages and over time.

**Figure 3:**
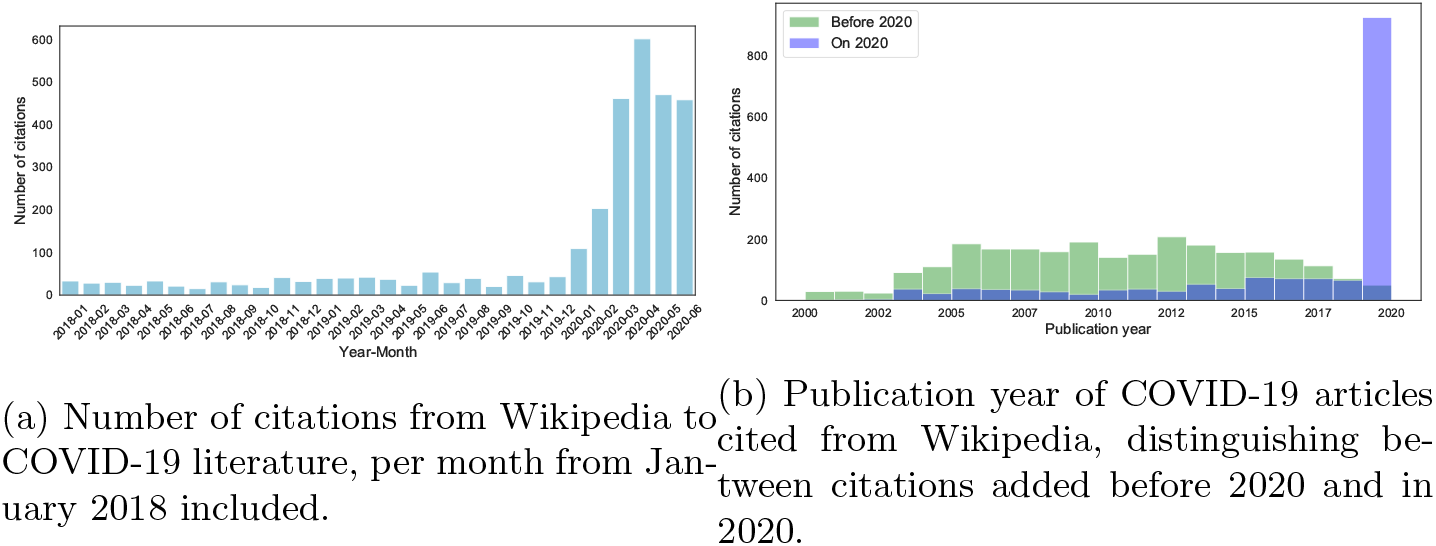
Timing of new citations from Wikipedia, and publication years of the articles they refer to. See Figure 7 for the full timeline.

Articles in English have a 2.0% chance of being cited from Wikipedia, while articles in other languages only a 0.24% chance. To be sure, the whole corpus is English dominated, as we discussed above. This might be an artefact of the coverage of the data sources, as well as the way the corpus was assembled. The coverage of articles over time is instead given in Figure 4, starting from 2003 when the first surge of publications happens due to SARS. We can appreciate that the coverage seems to be uneven, and less pronounced for the past few years (2017-2020), yet this needs to be considered in view of the high growth of publications in 2020. Hence, while 2020 is a relatively low-coverage year (1.2%), it is already the year with the most publications cited from Wikipedia in absolute number (Figure 3b).

**Figure 4:**
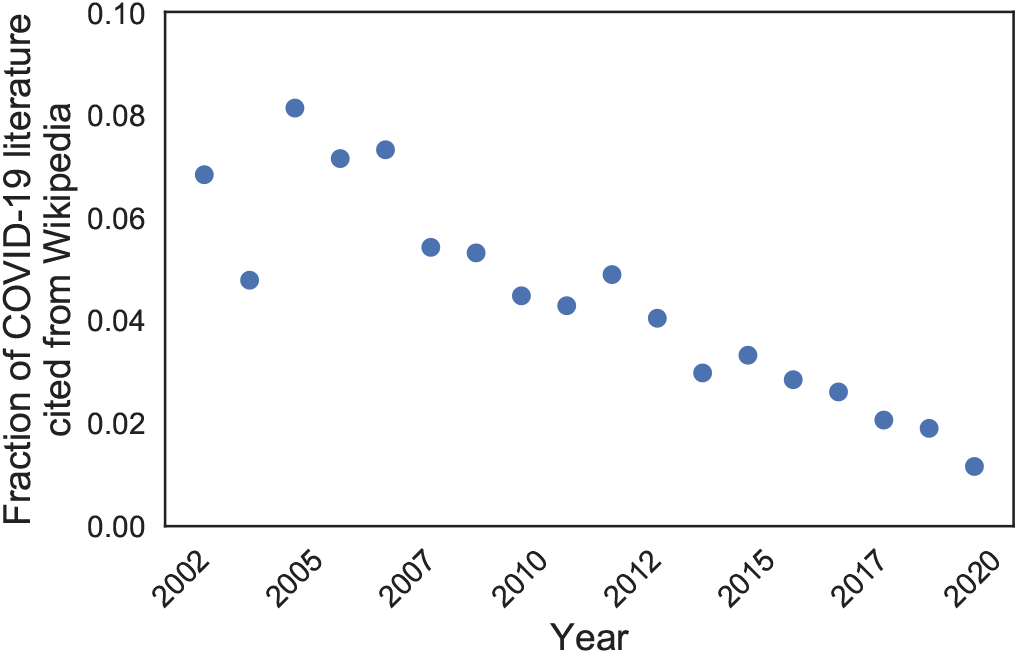
Fraction of COVID-19-related articles cited from Wikipedia per year, from 2003 included.

Citation distributions are skewed in Wikipedia as they are in science more generally. Some articles receive a high number of citations from Wikipedia and some Wikipedia articles make a high number of citations to COVID-19-related literature. Table 2 lists the top 20 Wikipedia articles by number of citations to COVID-19-related research. These articles, largely in English, primarily focus on the recent pandemic and coronaviruses/viruses from a virology perspective, as already highlighted in a study by the Wikimedia Foundation [24]. Table 3 reports instead the top 20 journal articles cited from Wikipedia. These also follow a similar pattern: articles published before 2020 focus on virology and are made of a high proportion of review articles. Articles published in 2020, instead, have a focus on the ongoing pandemic, its origins, as well as its epidemiological and public health aspects. As we see next, this strongly aligns with the general trends of COVID-19-related research over time.

In order to discuss research trends in our CORD-19-related corpus at a higher level of granularity, we grouped the 15 topics from the LDA topic model into five *general topics* and labelled them as follows:

- **Coronaviruses**: topics 5, 8; this general topic includes research explicitly on coronaviruses (COVID-19, SARS, MERS) from a variety of per-spectives (virology, epidemiology, intensive care, historical unfolding of outbreaks).
- **Epidemics**: topics 9, 11, 12; research on epidemiology, including modelling the transmission and spread of pathogens.
- **Public health**: topics 0, 1, 10; research on global health issues, healthcare.
- **Molecular biology and immunology**: topics 2, 4, 6; research on the genetics and biology of viruses, vaccines, drugs, therapies.
- **Clinical medicine**: topics 3, 7, 13, 14; research on intensive care, hospitalization and clinical trials.

The grouping is informed by agglomerative clustering based on the Jensen-Shannon distance between topic-word distributions (Figure 11). To be sure, the labelling is a simplification of the actual publication contents. It is also worth considering that topics overlap substantially. The COVID-19 research corpus is dominated by literature on coronaviruses, public health and epidemics, largely due to 2020 publications. COVID-19-related research did not accumulate uniformly over time. We plot the relative (yearly mean, Figure 9a) and absolute (yearly sum, Figure 9b) general topic intensity. From these plots, we confirm the periodisation of COVID-19-related research as connected to known outbreaks. Outbreaks generate a shift in the attention of the research community, which is apparent when we consider the relative general topic intensity over time in Figure 9a. The 2003 SARS outbreak generated a shift associated with a raise of publications on coronaviruses and on the management of epidemic outbreaks (public health, epidemiology). A similar shift is again happening, at a much larger scale, during the current COVID-19 pandemic. When we consider the absolute general topic intensity, which can be interpreted as the number of articles on a given topic (Figure 9b), we can appreciate how scientists are mostly focusing on topics related to public health, epidemics and coronaviruses (COVID-19) during these first months of the current pandemic.

### 4.1 RQ1: Wikipedia coverage of COVID-19-related research

We address here our first research question: *Is the literature cited from Wikipedia representative of the broader topics discussed in COVID-19-related research?* We start by comparing the general topic coverage of articles cited from Wikipedia with those which are not. In Figure 5, three plots are provided: the general topic intensity of articles published before 2020 (Figure 5a), in 2020 (Figure 5b) and overall (Figure 5c). The general topic intensity is averaged and 95% confidence intervals are provided. From Figure 5c we can see that Wikipedia seems to cover COVID-19-related research well. The general topics on immunology, molecular biology and epidemics seem slightly over represented, where clinical medicine and public health are slightly under represented. A comparison between publications from 2020 and from before highlights further trends. In particular, in 2020 Wikipedia editors have focused more on recent literature on coronaviruses, thus directly related to COVID-19 and the current pandemic, and proportionally less on literature on public health, which is also dominating 2020 publications. The traditional slight over representation of immunology and molecular biology literature persists. Detailed Kruskal-Wallis H test statistics for significant differences [29] and Cohen’s d for their effect sizes [14] are provided in the SI (Figure 12 and Tables 4, 5, 6). While distributions are significantly different for most general topics and periodisations, the effect sizes are often small. The coverage of COVID-19-related literature from Wikipedia appears therefore to be reasonably balanced from this first analysis, and to remain so in 2020. The topical differences we found, especially around coronaviruses and the current COVID-19 outbreak, might in part be explained by the criterion of notability which led to the creation or expansion of Wikipedia articles on the ongoing pandemic.^9^

**Figure 5:**
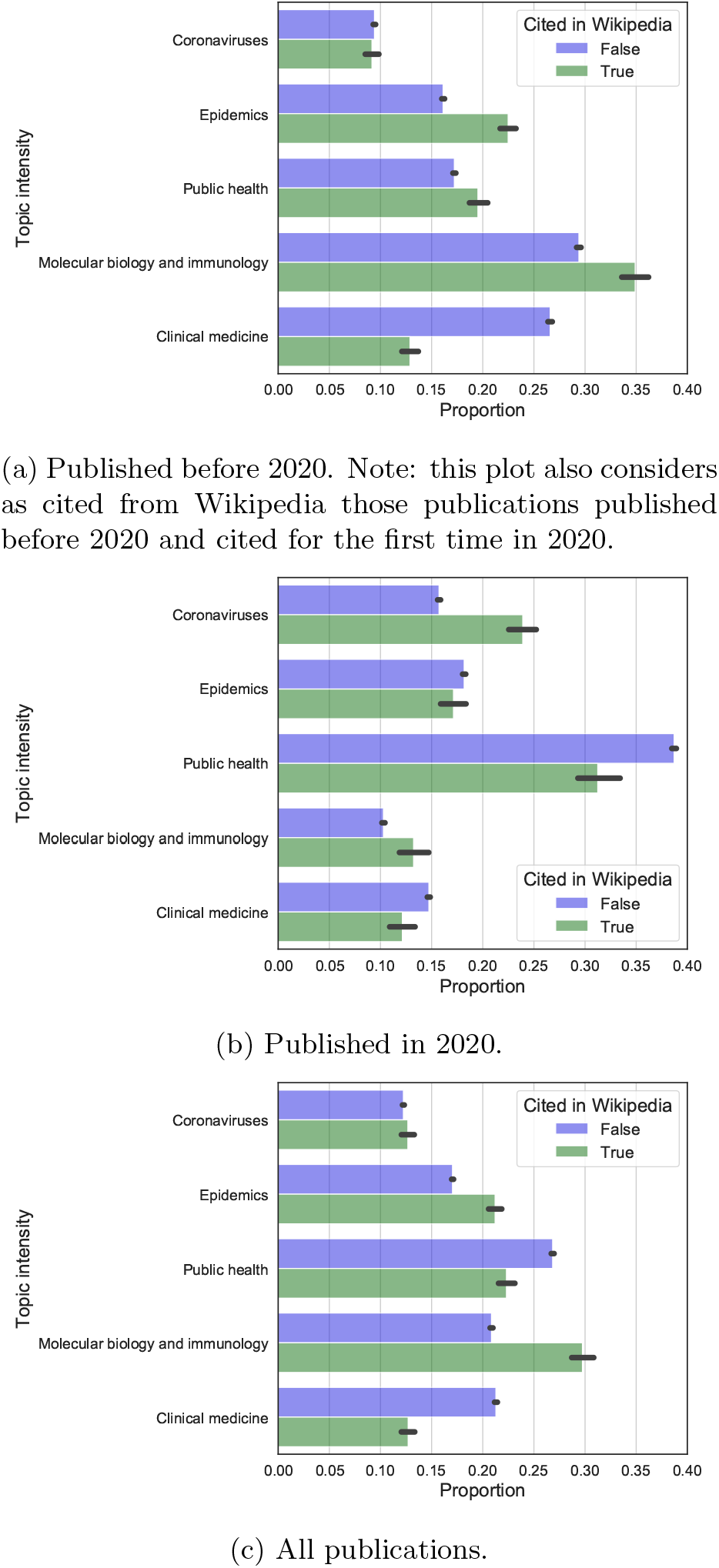
Average general topic intensity of COVID-19-related publications cited in Wikipeda (green) or not (blue). 95% bootstrapped confidence intervals are given. See Figure 12 and Tables 4, 5, 6 for significance tests and effect sizes.

A complementary way to address the same research question is to investigate Wikipedia’s coverage of publication clusters. We consider here both SPECTER k-means clusters and bibliographic network clusters. While we use all 20 SPECTER clusters, we limit ourselves to the top-n network clusters which are necessary in order to cover at least 50% of the nodes in the network. In this way, we consider 41 clusters for the citation network, all of size above 300. In Figure 6 we plot the % of articles cited from Wikipedia per cluster, and the clusters size in number of publications they contain. There is no apparent size effect in either of the two clustering solutions.

**Figure 6:**
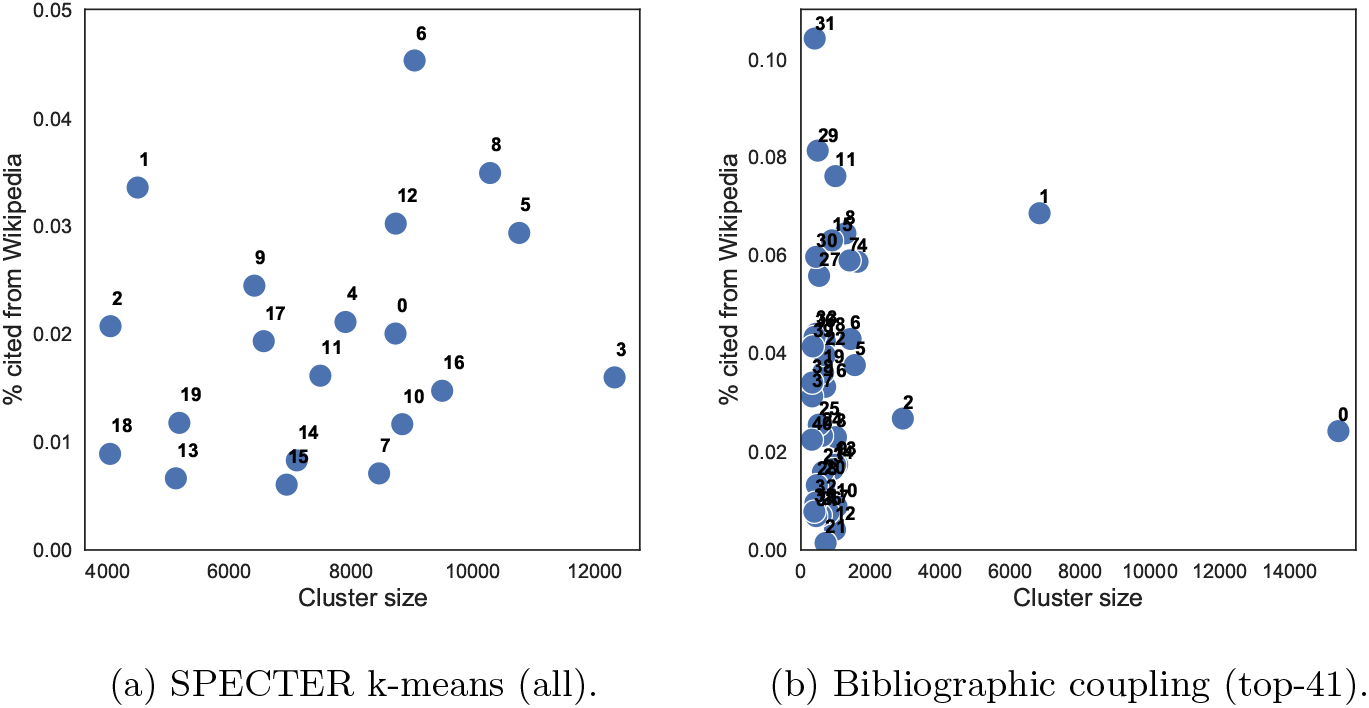
Proportion of articles cited from Wikipedia (y axis) per cluster size (number of articles in the cluster, x axis).

When we characterise clusters using general topic intensities, some clear patterns emerge. Starting with SPECTER k-means clusters, the most cited clusters are number 6 and 8 (main macrotopics: molecular biology) and 5 (main macrotopics: coronaviruses and public health, especially focusing on COVID-19 characteristics, detection and treatment). The least cited clusters include number 18 (containing pre-prints) and 13 (focused on the social sciences, and especially economics, e.g., from SSRN journals). Considering citation network clusters, the largest but not most cited are number 0 (containing 2020 research on COVID-19) and 1 (with publications on molecular biology and immunology). The other clusters are smaller and hence more specialized. The reader can explore all clusters using the accompanying repository.

We have seen so far that Wikipedia relies on a reasonably representative sample of COVID-19-related literature, when assessed using topic models. During 2020, the main effort of editors has focused on catching-up with abundant new research (and some backlog) on the ongoing pandemic and, to a lower extent, on public health and epidemiology literature. When assessing coverage using different clustering methods, we do not find a size effect by which larger clusters are proportionally more cited from Wikipedia. Yet we also find that, in particular with citation network clusters, smaller clusters can be either highly or lowly cited from Wikipedia on average. Lastly, we find an under representation of pre-print and social science research. Despite this overall encouraging result, differences in coverage persist. In the next section, we further assess whether these differences can be explained away by considering article-level measures of impact.

### 4.2 RQ2: Predictors of citations from Wikipedia

In this section, we address our second research question: *Is Wikipedia citing COVID-19-related research during the pandemic following the same criteria adopted before and in general?* We use regression analysis in two forms: a logistic regression to model if a paper is cited from Wikipedia or not, and a linear regression to model the number of citations a paper receives from Wikipedia. While the former model captures the suitability of an article to provide encyclopedic evidence, the latter captures its relevance to multiple Wikipedia articles.

#### Dependent variables

*Wikipedia citation counts* for each article are taken from Altmetric. If this count is of 1 or more, an article is considered as cited from Wikipedia. We consider citation counts from Altmetric at the time of the data collection for this study. We focus on the articles with a match from Dimensions, and consider an article to have zero citations from Wikipedia if it is not found in the Altmetric database.

#### Independent variables

We focus our study on three groups of independent variables at the article level capturing impact, topic and timing respectively. Previous studies have shown how literature cited from Wikipedia tends to be published in prestigious journals and available in open access [41, 59, 6]. We are interested to assess some of these known patterns for COVID-19-related research, to complement them by considering citation counts and the topics discussed in the literature, and eventually to understand whether there has been any change in 2020.

Article-level variables include citation counts from Dimensions and a variety of altmetric indicators [51] which have been found to correlate with later citation impact of COVID-19 research [28]. Altmetrics include the number of: Mendeley readers, Twitter interactions (unique users), Facebook shares, mentions in news and blog posts (summed due to their high correlation), mentions in policy documents; the expert ratio in user engagement^10^. We also include the top-20 publication venues by number of articles in the corpus using dummy coding, taking as reference level a generic category ‘other’ which includes articles from all other venues. It is worth clarifying that article-level variables were also calculated at the time of the data collection for this study. This might seem counter-intuitive, especially for the classification task, as one might prefer to calculate variables at the time when an article was first cited from Wikipedia. We argue that this is not the case, since Wikipedia can always be edited and citations removed as easily as added. As a consequence, a citation from Wikipedia (or its absence) is a continued rather than a discrete action, justifying calculating all counts at the same time for all articles in the corpus.

Topic-level variables capture the topics discussed in the articles, as well as their relative importance in terms of size (size-effects). They include the macrotopic intensities for each article, the size of the SPECTER cluster an article belongs to, and the size of its bibliographic coupling network cluster (for the 41 largest clusters with more than 300 articles each, setting it to zero for articles belonging to other clusters. In this way, the variable accounts for both size and thresholding effects). Cluster identities for both SPECTRE and citation network clusters were also tested but did not contribute significantly to the models. Several other measures were considered, such as the semantic centrality of an article to its cluster centroid (SPECTER k-means) and network centralities, but since these all strongly correlate to size indicators, they were discarded to avoid multicollinearity.

Lastly, we include the year of publication using dummy coding and 2020 as reference level. Several other variables were tested. The proposed selection removes highly correlated variables while preserving the information required by the research question. The Pearson’s correlations for the selected transformed variables are shown in Figure 10. More details, along with a full profiling of variables, are provided in the accompanying repository.

#### Model

We consider two models: a Logistic model on being cited from Wikipedia (1) or not (0) and an Ordinary Least Squares (OLS) model on citation counts from Wikipedia. Both models use the same set of independent variables and transformations described in Table 1.

**Table 1:** Regression variables, their description, typology and transformations. *In*(*x* + 1) means one is added to the value and then the natural logarithm is taken.

| Variable | Description | Type | Transformations |
| --- | --- | --- | --- |
| *in_wikipedia* | Whether an article is cited from Wikipedia (1) or not (0) | Indicator |  |
| *n_cit_w* | Number of citations from Wikipedia | Numeric | $\ln(x+1)$ |
| *publication_year* | Publication year of the article | Categorical |  |
| *times_cited* | Number of citations (Dimensions) | Numeric | $\ln(x+1)$ |
| *counts_mendeley* | Number of Mendeley readers (Altmetric) | Numeric | $\ln(x+1)$ |
| *counts_policy* | Number of mentions from policy documents (Altmetric) | Numeric | $\ln(x+1)$ |
| *counts_twitter_unique* | Number of engagements from unique Twitter users (Altmetric) | Numeric | $\ln(x+1)$ |
| *counts_blogs_news* | Number of mentions in news and blogs (Altmetric) | Numeric | $\ln(x+1)$ |
| *counts_facebook* | Number of mentions in Facebook (Altmetric) | Numeric | $\ln(x+1)$ |
| *expert_ratio* | Ratio of engagements from experts (Altmetric) | Numeric (0 to 1) |  |
| *top_journal* | Journal | Categorical |  |
| *tm_coronaviruses* | Topic intensity: Coronaviruses | Numeric (0 to 1) |  |
| *tm_epidemics* | Topic intensity: Epidemics | Numeric (0 to 1) |  |
| *tm_ph* | Topic intensity: Public health | Numeric (0 to 1) |  |
| *tm_mbi* | Topic intensity: Molecular biology and immunology | Numeric (0 to 1) |  |
| *tm_clinical_medicine* | Topic intensity: Clinical medicine | Numeric (0 to 1) |  |
| *spectre_cluster_size* | Size of SPECTRE cluster the article belongs to | Numeric | $\ln(x+1)$ |
| *network_cluster_size* | Size of bib. coupling cluster the article belongs to | Numeric | $\ln(x+1)$ |

All count variables are transformed by adding one and taking the natural logarithm, while the remaining variables are either indicators or range between 0 and 1 (such as general topic intensities, beginning with a *tm*_ prefix; e.g., *tm_ph* is ‘public health’). OLS models including log transform and the addition of 1 for count variables such as citation counts, have been found to perform well in practice when compared to more involved alternatives [61, 60]. Furthermore, all missing values were set to zero, except for the publication year, venue (journal) and general topic intensities since removing rows with missing values yielded comparable results.

#### Discussion

We discuss results for three models: two Logistic regression models one on articles published and first cited up to and including in 2020, and one on articles published and first cited up to an including 2019. The 2019 model only considers articles published in 2019 or earlier and cited for the first time from Wikipedia in 2019 or earlier, or articles never cited from Wikipedia, discarding articles published in 2020 or cited from Wikipedia in 2020 irrespective of their publication time. We also discuss an OLS model predicting (the log of) citation counts including all data up to and including 2020. We do not discuss a 2019 OLS model since it would require Wikipedia citation counts calculated at the end of 2019, which were not available to us. Regression tables for these three models are provided in the SI, Section 5, while Figure 13 shows the distribution of some variables distinguishing between articles cited from Wikipedia or not. Logistic regression tables provide marginal effects, while the OLS table provides the usual coefficients. The actual number of datapoints used to fit each model, after removing those which contained any null value, is given in the regression tables.

Considering the Logistic models first, we can show some significant effects.^11^ First of all, the year of publication is mostly negatively correlated with being cited from Wikipedia, compared with the reference category 2020. This seems largely due to publication size-effects, since the fraction of 2020 articles cited from Wikipedia is quite low (see Figure 4). The 2019 model indeed shows positive correlations for all years when compared to the reference category 2019, and indeed 2019 is the year with lowest coverage since 2000. Secondly, some of the most popular venues are positively correlated with citations from Wikipedia, when compared to an ‘other’ category (which includes all venues except the top 20). In the 2020 model, these venues include mega-journals (Nature, Science) and specialized journals (The Lancet, BMJ). Negative correlations occur for pre-print servers (medRxiv and bioRxiv in particular).

When we consider indicators of impact, we see a significant positive effect for citation counts, Mendeley readers, Twitter, news and blogs mentions; we see instead no effect for policy document mentions and Facebook engagements. This is consistent in the 2019 model, except for Facebook having a positive effect and Twitter a lack of correlation. This result, on the one hand, highlights the importance of academic indicators of impact such as citations, and on the other hand suggests the possible complementarity of altmetrics in this respect. Since certain altmetrics can accumulate more rapidly than citations [16], they could complement them effectively when needed [28]. Furthermore, the expert ratio in altmetrics engagement is negatively correlated with being cited from Wikipedia in 2020. This might be due to the high altmetrics engagement with COVID-19 research in 2020, but it could also hint at the possibility that social media impact need not be driven by experts in order to be correlated with scientific impact. We can further see how cluster size-effects are not or very marginally correlated with being cited from Wikipedia.

Lastly, *general topic intensities are never correlated with being cited from Wikipedia in either model,* underlining that Wikipedia appears to be proportionally representing all COVID-19-related research and that residual topical differences in coverage are due to article-level effects.

The 2020 OLS model largely confirms these results, except that mentions in policy documents and Facebook engagements become positively correlated with the number of citations from Wikipedia. It is important to underline that, for all these results, there is no attempt to establish causality. For example, the positive correlation between the number of Wikipedia articles citing a scientific article and the number of policy documents mentioning it, might be due to policy document editors using Wikipedia, Wikipedia editors using policy documents, both or neither. The fact is, more simply, that some articles are picked up by both.

## 5 Conclusion

The results of this study provide some reassuring evidence. It appears that Wikipedia’s editors are well-able to keep track of COVID-19-related research. Of 141,783 articles in our corpus, 3083 (~2%) are cited from Wikipedia: a share comparable to what found in previous studies. Wikipedia editors are relying on scientific results representative of the several topics included in a large corpus of COVID-19-related research. They have been effectively able to cope with new, rapidly-growing literature. The minor discrepancies in coverage that persist, with slightly more Wikipedia-cited articles on topics such as molecular biology and immunology and slightly fewer on clinical medicine and public health, are fully explained away by article-level effects. Wikipedia editors rely on impactful and visible research, as evidenced by largely positive citation and altmetrics correlations. Importantly, Wikipedia editors also appear to be following the same inclusion standards in 2020 as before: in general, they rely on specialized and highly-cited results from reputed journals, avoiding e.g., pre-prints.

The main limitation of this study is that it is purely observational, and thus does not explain why some articles are cited from Wikipedia or not. While in order to assess the coverage of COVID-19-related research from Wikipedia this is of secondary importance, it remains relevant when attempting to predict and explain it. A second limitation is that this study is based on citations from Wikipedia to scientific publications, and no Wikipedia content analysis is performed. Citations to scientific literature, while informative, do not completely address the interrelated questions of Wikipedia’s knowledge representativeness and reliability. Therefore, some directions for future work include comparing Wikipedia coverage with expert COVID-19 review articles, as well as studying Wikipedia edit and discussion history in order to assess editor motivations. Another interesting direction for future work is the assessment of all Wikipedia citations to any source from COVID-19 Wikipedia pages, since here we only focused on the fraction directed at COVID-19-related scientific articles. Lastly, future work can address the engagement of Wikipedia users with cited COVID-19-related sources.

Wikipedia is a fundamental source of free knowledge, open to all. The capacity of its editor community to quickly respond to a crisis and provide high-quality contents is, therefore, critical. Our results here are encouraging in this respect.

## Data and code availability

All the analyses can be replicated using code and following the instructions given in the accompanying repository: https://github.com/Giovanni1085/covid-19wikipedia. The preparation of the data follows the steps detailed in this repository instead: https://github.com/CWTSLeiden/cwts_covid [15]. Analyses based on Altmetric and Dimensions data require access to these services.

## Acknowledgements

Digital Science kindly provided access to Altmetric and Dimensions data.

## SI

### Topics

Refer to Figures 8 and 9 for topic intensities over time. See Figure 11 for the topic clustering. The topic label is given next to the topic number, for reference.

- **Topic #0, Public health**: “method”, “system”, “use”, “drug”, “application”, “approach”, “image”, “design”, “test”, “develop”, “technology”, “provide”, “technique”, “new”, “tool”, “potential”, “base”, “device”, “allow”, “result”.
- **Topic #1, Public health**: “health”, “pandemic”, “covid-19”, “COVID-19”, “public”, “country”, “outbreak”, “social”, “care”, “covid-19_pandemic”, “measure”, “policy”, “people”, “public_health”, “Health”, “impact”, “response”, “risk”, “medical”, “need”.
- **Topic #2, Molecular biology and immunology**: “cell”, “infection”, “response”, “mouse”, “immune”, “expression”, “lung”, “induce”, “disease”, “cat”, “role”, “tissue”, “system”, “increase”, “level”, “receptor”, “study”, “gene”, “cytokine”, “human”.
- **Topic #3, Clinical medicine**: “group”, “patient”, “day”, “study”, “year”, “result”, “rate”, “age”, “method”, “compare”, “conclusion”, “total”, “time”, “period”, “mean”, “respectively”, “high”, “month”, “significantly”.
- **Topic #4, Molecular biology and immunology**: “protein”, “virus”, “cell”, “rna”, “viral”, “coronavirus”, “activity”, “replication”, “gene”, “antiviral”, “study”, “human”, “membrane”, “domain”, “binding”, “structure”, “sequence”, “target”, “infection”, “inhibitor”.
- **Topic #5, Coronaviruses**: “respiratory”, “infection”, “acute”, “virus”, “syndrome”, “SARS”, “severe”, “respiratory_syndrome”, “severe_acute”, “influenza”, “child”, “case”, “patient”, “viral”, “acute respiratory syndrome”, “cause”, “coronavirus”, “clinical”, “sars”, “pneumonia”.
- **Topic #6, Molecular biology and immunology**: “virus”, “antibody”, “strain”, “sample”, “detect”, “sequence”, “assay”, “isolate”, “coronavirus”, “detection”, “test”, “gene”, “calf”, “result”, “serum”, “positive”, “analysis”, “study”, “bovine”, “ibv”.
- **Topic #7, Clinical medicine**: “patient”, “surgery”, “laparoscopic”, “surgical”, “procedure”, “cancer”, “complication”, “perform”, “technique”, “undergo”, “postoperative”, “case”, “tumor”, “result”, “method”, “repair”, “time”, “patient_undergo”, “resection”, “hernia”.
- **Topic #8, Coronaviruses**: “covid-19”, “COVID-19”, “sars-cov-2”, “coronavirus”, “case”, “disease”, “patient”, “2019”, “2020”, “infection”, “severe”, “clinical”, “China”, “novel”, “confirm”, “coronavirus_disease”, “re-port”, “symptom”, “novel_coronavirus”, “Wuhan”.
- **Topic #9, Epidemics**: “model”, “datum”, “number”, “analysis”, “epidemic”, “case”, “time”, “network”, “study”, “different”, “result”, “rate”, “dynamic”, “base”, “paper”, “estimate”, “propose”, “population”, “spread”, “individual”.
- **Topic #10, Public health**: “study”, “review”, “trial”, “include”, “clinical”, “treatment”, “search”, “evidence”, “literature”, “result”, “datum”, “intervention”, “quality”, “report”, “systematic”, “use”, “outcome”, “method”, “research”, “article”.
- **Topic #11, Epidemics**: “disease”, “vaccine”, “infectious”, “human”, “review”, “virus”, “new”, “infectious_disease”, “emerge”, “development”, “animal”, “infection”, “pathogen”, “recent”, “potential”, “cause”, “vaccination”, “infectious diseases”, “outbreak”, “include”.
- **Topic #12, Epidemics**: “risk”, “factor”, “associate”, “associated with”, “mortality”, “high”, “analysis”, “increase”, “study”, “95_ci”, “risk_factor”, “death”, “age”, “patient”, “rate”, “ratio”, “outcome”, “regression”.
- **Topic #13, Clinical medicine**: “effect”, “increase”, “group”, “study”, “level”, “concentration”, “control”, “blood”, “change”, “pressure”, “result”, “high”, “low”, “decrease”, “compare”, “measure”, “temperature”, “significantly”, “weight”, “reduce”.
- **Topic #14, Clinical medicine**: “patient”, “treatment”, “clinical”, “acute”, “lung”, “therapy”, “chest”, “aneurysm”, “outcome”, “treat”, “ventilation”, “care”, “case”, “artery”, “stroke”, “failure”, “lesion”, “pulmonary”, “diagnosis”.

### Extra tables and figures

**Table 2:** Top-20 citing Wikipedia articles.

| # citations | Wikipedia id | Wikipedia article title | Lang |
| --- | --- | --- | --- |
| 62 | 201983 | Coronavirus | en |
| 59 | 62786585 | Severe acute respiratory syndrome coronavirus 2 | en |
| 53 | 62750956 | 2019–20 Wuhan coronavirus outbreak | en |
| 49 | 63030231 | Coronavirus disease 2019 | en |
| 39 | 63676463 | 2019–20 coronavirus pandemic | en |
| 36 | 63430824 | COVID-19 drug repurposing research | en |
| 33 | 63895130 | Paediatric multisystem inflammatory syndrome | en |
| 32 | 211547 | Severe acute respiratory syndrome-related coro... | en |
| 31 | 63319438 | COVID-19 vaccine | en |
| 30 | 63435931 | COVID-19 drug development | en |
| 34 | 39532251 | Middle East respiratory syndrome | en |
| 28 | 19572217 | Influenza | en |
| 25 | 63204759 | COVID-19 testing | en |
| 23 | 2717089 | Angiotensin-converting enzyme 2 | en |
| 24 | 22693252 | Feline coronavirus | en |
| 22 | 64144585 | Management of COVID-19 | en |
| 22 | 4354646 | Emergent virus | en |
| 21 | 10849236 | Antibody-dependent enhancement | en |
| 20 | 196741 | Severe acute respiratory syndrome | en |
| 20 | 64144627 | Prognosis of COVID-19 | en |

**Table 3:**
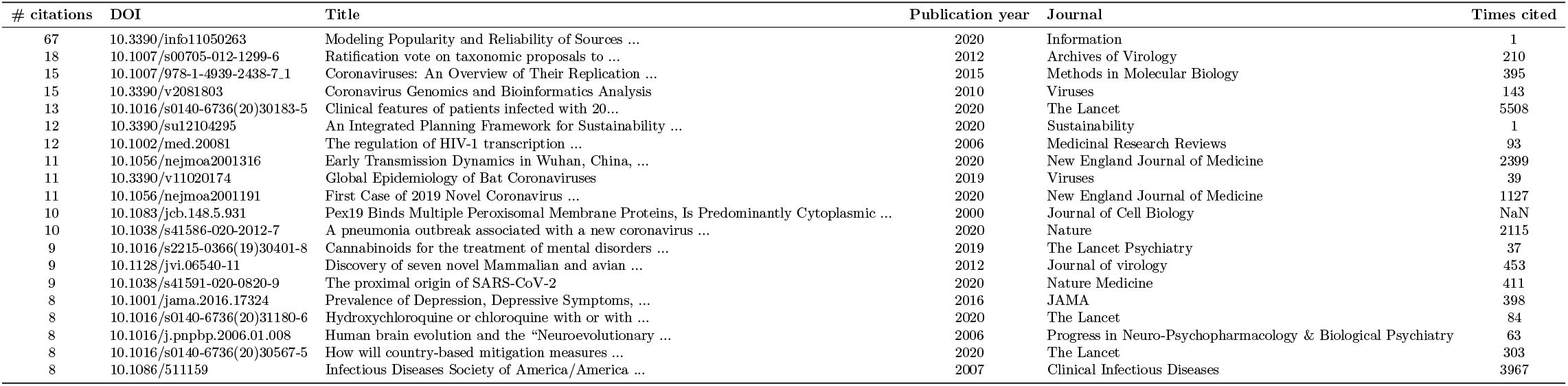
Top-20 cited journal articles. The first column gives the number of distinct citing Wikipedia articles, while the last one gives the number of citations to these articles from the scientific literature (data from Dimensions).

### Regression tables

| **Model:** | Logistic regression 2020 |  |  |  |  |  |
| --- | --- | --- | --- | --- | --- | --- |
| **Method:** | Marginal effects (Logistic regression) |  |  |  |  |  |
| **No. Observations:** | 130,864 |  |  |  |  |  |
| **Pseudo R-squ.:** | 0.2790 |  |  |  |  |  |
| variable | dx/dy coef | std err | z | P> \|z\| | [0.025 | 0.975] |
| C(publication_year, Treatment(2020))[T.2000.0] | -0.0035 | 0.008 | -0.457 | 0.648 | -0.019 | 0.012 |
| C(publication_year, Treatment(2020))[T.2001.0] | -0.0164 | 0.010 | -1.644 | 0.100 | -0.036 | 0.003 |
| C(publication_year, Treatment(2020))[T.2002.0] | -0.0018 | 0.006 | -0.317 | 0.751 | -0.013 | 0.009 |
| C(publication_year, Treatment(2020))[T.2003.0] | -0.0030 | 0.004 | -0.763 | 0.445 | -0.011 | 0.005 |
| C(publication_year, Treatment(2020))[T.2004.0] | 0.0004 | 0.003 | 0.127 | 0.899 | -0.005 | 0.006 |
| C(publication_year, Treatment(2020))[T.2005.0] | 0.0050 | 0.003 | 1.755 | 0.079 | -0.001 | 0.011 |
| C(publication_year, Treatment(2020))[T.2006.0] | 0.0036 | 0.003 | 1.303 | 0.192 | -0.002 | 0.009 |
| C(publication_year, Treatment(2020))[T.2007.0] | 0.0023 | 0.003 | 0.792 | 0.428 | -0.003 | 0.008 |
| C(publication_year, Treatment(2020))[T.2008.0] | 0.0034 | 0.002 | 1.357 | 0.175 | -0.002 | 0.008 |
| C(publication_year, Treatment(2020))[T.2009.0] | -0.0010 | 0.003 | -0.388 | 0.698 | -0.006 | 0.004 |
| C(publication_year, Treatment(2020))[T.2010.0] | 0.0010 | 0.002 | 0.392 | 0.695 | -0.004 | 0.006 |
| C(publication_year, Treatment(2020))[T.2011.0] | -0.0031 | 0.002 | -1.260 | 0.208 | -0.008 | 0.002 |
| C(publication_year, Treatment(2020))[T.2012.0] | -0.0032 | 0.002 | -1.375 | 0.169 | -0.008 | 0.001 |
| C(publication_year, Treatment(2020))[T.2013.0] | -0.0080 | 0.002 | -3.458 | 0.001 | -0.012 | -0.003 |
| C(publication_year, Treatment(2020))[T.2014.0] | -0.0085 | 0.002 | -3.842 | 0.000 | -0.013 | -0.004 |
| C(publication_year, Treatment(2020))[T.2015.0] | -0.0123 | 0.002 | -5.528 | 0.000 | -0.017 | -0.008 |
| C(publication_year, Treatment(2020))[T.2016.0] | -0.0119 | 0.002 | -5.321 | 0.000 | -0.016 | -0.008 |
| C(publication_year, Treatment(2020))[T.2017.0] | -0.0116 | 0.002 | -5.191 | 0.000 | -0.016 | -0.007 |
| C(publication_year, Treatment(2020))[T.2018.0] | -0.0112 | 0.002 | -5.001 | 0.000 | -0.016 | -0.007 |
| C(publication_year, Treatment(2020))[T.2019.0] | -0.0067 | 0.002 | -2.919 | 0.004 | -0.011 | -0.002 |
| C(top_j, Treatment('OTHER'))[T.Arch_Virol] | 0.0031 | 0.006 | 0.528 | 0.598 | -0.008 | 0.015 |
| C(top_j, Treatment('OTHER'))[T.ChemRxiv] | -0.0010 | 0.011 | -0.089 | 0.929 | -0.023 | 0.021 |
| C(top_j, Treatment('OTHER'))[T.Emerg_Infect_Dis] | -0.0001 | 0.003 | -0.051 | 0.960 | -0.006 | 0.005 |
| C(top_j, Treatment('OTHER'))[T.JAMA] | -0.0222 | 0.004 | -5.200 | 0.000 | -0.031 | -0.014 |
| C(top_j, Treatment('OTHER'))[T.JMIR_Preprints] | -0.2486 | 10.693 | -0.023 | 0.981 | -21.207 | 20.710 |
| C(top_j, Treatment('OTHER'))[T.Journal_of_virology] | -0.0013 | 0.003 | -0.422 | 0.673 | -0.007 | 0.005 |
| C(top_j, Treatment('OTHER'))[T.Nature] | 0.0105 | 0.003 | 3.614 | 0.000 | 0.005 | 0.016 |
| C(top_j, Treatment('OTHER'))[T.PLoS_One] | -0.0034 | 0.003 | -1.255 | 0.210 | -0.009 | 0.002 |
| C(top_j, Treatment('OTHER'))[T.Research_Square] | -0.3905 | 235.195 | -0.002 | 0.999 | -461.365 | 460.584 |
| C(top_j, Treatment('OTHER'))[T.SSRN_Electronic_Journal] | -0.0173 | 0.008 | -2.208 | 0.027 | -0.033 | -0.002 |
| C(top_j, Treatment('OTHER'))[T.Sci_Rep] | -0.0071 | 0.005 | -1.375 | 0.169 | -0.017 | 0.003 |
| C(top_j, Treatment('OTHER'))[T.Science] | 0.0083 | 0.004 | 2.202 | 0.028 | 0.001 | 0.016 |
| C(top_j, Treatment('OTHER'))[T.Surgical_Endoscopy] | -0.0003 | 0.007 | -0.039 | 0.969 | -0.014 | 0.013 |
| C(top_j, Treatment('OTHER'))[T.The_BMJ] | 0.0126 | 0.004 | 3.267 | 0.001 | 0.005 | 0.020 |
| C(top_j, Treatment('OTHER'))[T.The_Lancet] | 0.0129 | 0.003 | 4.645 | 0.000 | 0.007 | 0.018 |
| C(top_j, Treatment('OTHER'))[T.Vaccine] | -0.0118 | 0.006 | -1.909 | 0.056 | -0.024 | 0.000 |
| C(top_j, Treatment('OTHER'))[T.Virology] | -0.0013 | 0.004 | -0.308 | 0.758 | -0.010 | 0.007 |
| C(top_j, Treatment('OTHER'))[T.Viruses] | 0.0091 | 0.003 | 2.639 | 0.008 | 0.002 | 0.016 |
| C(top_j, Treatment('OTHER'))[T.bioRxiv] | -0.0073 | 0.004 | -1.824 | 0.068 | -0.015 | 0.001 |
| C(top_j, Treatment('OTHER'))[T.medRxiv] | -0.0489 | 0.008 | -6.260 | 0.000 | -0.064 | -0.034 |
| times_cited | 0.0078 | 0.000 | 19.909 | 0.000 | 0.007 | 0.009 |
| counts_mendeley | 0.0066 | 0.000 | 20.967 | 0.000 | 0.006 | 0.007 |
| counts_policy | -0.0013 | 0.001 | -1.319 | 0.187 | -0.003 | 0.001 |
| counts_twitter_unique | 0.0050 | 0.000 | 13.822 | 0.000 | 0.004 | 0.006 |
| counts_blogs_news | 0.0051 | 0.000 | 11.343 | 0.000 | 0.004 | 0.006 |
| counts_facebook | 0.0008 | 0.001 | 1.115 | 0.265 | -0.001 | 0.002 |
| expert_ratio | -0.0067 | 0.002 | -3.679 | 0.000 | -0.010 | -0.003 |
| tm_coronaviruses | 0.0138 | 0.018 | 0.760 | 0.447 | -0.022 | 0.050 |
| tm_epidemics | 0.0241 | 0.018 | 1.343 | 0.179 | -0.011 | 0.059 |
| tm_ph | 0.0197 | 0.018 | 1.086 | 0.278 | -0.016 | 0.055 |
| tm_mbi | 0.0219 | 0.018 | 1.226 | 0.220 | -0.013 | 0.057 |
| tm_clinical_medicine | -0.0064 | 0.018 | -0.362 | 0.717 | -0.041 | 0.028 |
| spectre_cluster_size | 0.0022 | 0.001 | 1.483 | 0.138 | -0.001 | 0.005 |
| network_cluster_size | -2.14e-05 | 0.000 | -0.132 | 0.895 | -0.000 | 0.000 |

| variable | dx/dy coef | std err | z | P> \|z\| | [0.025 | 0.975] |
| --- | --- | --- | --- | --- | --- | --- |
| C(publication_year, Treatment(2019))[T.2000.0] | 0.0326 | 0.010 | 3.389 | 0.001 | 0.014 | 0.051 |
| C(publication_year, Treatment(2019))[T.2001.0] | 0.0190 | 0.012 | 1.615 | 0.106 | -0.004 | 0.042 |
| C(publication_year, Treatment(2019))[T.2002.0] | 0.0359 | 0.008 | 4.728 | 0.000 | 0.021 | 0.051 |
| C(publication_year, Treatment(2019))[T.2003.0] | 0.0322 | 0.006 | 5.487 | 0.000 | 0.021 | 0.044 |
| C(publication_year, Treatment(2019))[T.2004.0] | 0.0335 | 0.005 | 6.812 | 0.000 | 0.024 | 0.043 |
| C(publication_year, Treatment(2019))[T.2005.0] | 0.0415 | 0.005 | 8.825 | 0.000 | 0.032 | 0.051 |
| C(publication_year, Treatment(2019))[T.2006.0] | 0.0365 | 0.005 | 7.797 | 0.000 | 0.027 | 0.046 |
| C(publication_year, Treatment(2019))[T.2007.0] | 0.0371 | 0.005 | 7.913 | 0.000 | 0.028 | 0.046 |
| C(publication_year, Treatment(2019))[T.2008.0] | 0.0356 | 0.004 | 8.032 | 0.000 | 0.027 | 0.044 |
| C(publication_year, Treatment(2019))[T.2009.0] | 0.0297 | 0.004 | 6.776 | 0.000 | 0.021 | 0.038 |
| C(publication_year, Treatment(2019))[T.2010.0] | 0.0290 | 0.004 | 6.644 | 0.000 | 0.020 | 0.038 |
| C(publication_year, Treatment(2019))[T.2011.0] | 0.0247 | 0.004 | 5.730 | 0.000 | 0.016 | 0.033 |
| C(publication_year, Treatment(2019))[T.2012.0] | 0.0262 | 0.004 | 6.337 | 0.000 | 0.018 | 0.034 |
| C(publication_year, Treatment(2019))[T.2013.0] | 0.0201 | 0.004 | 4.874 | 0.000 | 0.012 | 0.028 |
| C(publication_year, Treatment(2019))[T.2014.0] | 0.0195 | 0.004 | 4.813 | 0.000 | 0.012 | 0.027 |
| C(publication_year, Treatment(2019))[T.2015.0] | 0.0129 | 0.004 | 3.189 | 0.001 | 0.005 | 0.021 |
| C(publication_year, Treatment(2019))[T.2016.0] | 0.0100 | 0.004 | 2.418 | 0.016 | 0.002 | 0.018 |
| C(publication_year, Treatment(2019))[T.2017.0] | 0.0110 | 0.004 | 2.668 | 0.008 | 0.003 | 0.019 |
| C(publication_year, Treatment(2019))[T.2018.0] | 0.0085 | 0.004 | 2.005 | 0.045 | 0.000 | 0.017 |
| C(top_j, Treatment('OTHER'))[T.Arch_Virol] | 0.0043 | 0.008 | 0.549 | 0.583 | -0.011 | 0.020 |
| C(top_j, Treatment('OTHER'))[T.Emerg_Infect_Dis] | 0.0022 | 0.004 | 0.574 | 0.566 | -0.005 | 0.010 |
| C(top_j, Treatment('OTHER'))[T.JAMA] | 0.0067 | 0.007 | 1.029 | 0.303 | -0.006 | 0.020 |
| C(top_j, Treatment('OTHER'))[T.Journal_of_virology] | -0.0006 | 0.004 | -0.153 | 0.879 | -0.008 | 0.007 |
| C(top_j, Treatment('OTHER'))[T.Nature] | 0.0148 | 0.005 | 2.861 | 0.004 | 0.005 | 0.025 |
| C(top_j, Treatment('OTHER'))[T.PLoS_One] | -0.0070 | 0.004 | -1.926 | 0.054 | -0.014 | 0.000 |
| C(top_j, Treatment('OTHER'))[T.Sci_Rep] | 0.0002 | 0.007 | 0.027 | 0.978 | -0.013 | 0.013 |
| C(top_j, Treatment('OTHER'))[T.Science] | -0.0008 | 0.009 | -0.096 | 0.923 | -0.018 | 0.016 |
| C(top_j, Treatment('OTHER'))[T.Surgical_endoscopy] | 0.0037 | 0.008 | 0.477 | 0.633 | -0.011 | 0.019 |
| C(top_j, Treatment('OTHER'))[T.The_Lancet] | 0.0109 | 0.005 | 2.136 | 0.033 | 0.001 | 0.021 |
| C(top_j, Treatment('OTHER'))[T.Vaccine] | -0.0220 | 0.010 | -2.291 | 0.022 | -0.041 | -0.003 |
| C(top_j, Treatment('OTHER'))[T.Virology] | -0.0048 | 0.006 | -0.826 | 0.409 | -0.016 | 0.007 |
| C(top_j, Treatment('OTHER'))[T.Viruses] | 0.0078 | 0.005 | 1.562 | 0.118 | -0.002 | 0.018 |
| C(top_j, Treatment('OTHER'))[T.bioRxiv] | -0.9986 | 4.64e+08 | -2.15e-09 | 1.000 | -9.1e+08 | 9.1e+08 |
| times_cited | 0.0027 | 0.001 | 4.136 | 0.000 | 0.001 | 0.004 |
| counts_mendeley | 0.0164 | 0.001 | 22.952 | 0.000 | 0.015 | 0.018 |
| counts_policy | -0.0020 | 0.001 | -1.377 | 0.168 | -0.005 | 0.001 |
| counts_twitter_unique | 0.0006 | 0.001 | 0.915 | 0.360 | -0.001 | 0.002 |
| counts_blogs_news | 0.0051 | 0.001 | 6.915 | 0.000 | 0.004 | 0.007 |
| counts_facebook | 0.0044 | 0.001 | 3.950 | 0.000 | 0.002 | 0.007 |
| expert_ratio | -0.0010 | 0.002 | -0.408 | 0.683 | -0.006 | 0.004 |
| tm_coronaviruses | -0.0251 | 0.034 | -0.729 | 0.466 | -0.093 | 0.042 |
| tm_epidemics | -0.0172 | 0.034 | -0.505 | 0.613 | -0.084 | 0.049 |
| tm_ph | -0.0118 | 0.034 | -0.346 | 0.729 | -0.079 | 0.055 |
| tm_mbi | -0.0091 | 0.034 | -0.267 | 0.789 | -0.076 | 0.057 |
| tm_clinical_medicine | -0.0356 | 0.034 | -1.056 | 0.291 | -0.102 | 0.031 |
| spectre_cluster_size | 0.0041 | 0.002 | 1.947 | 0.052 | -2.79e-05 | 0.008 |
| network_cluster_size | -0.0009 | 0.000 | -3.371 | 0.001 | -0.001 | -0.000 |

| variable | coef | std err | t | P>\|t\| | [0.025 | 0.975] |
| --- | --- | --- | --- | --- | --- | --- |
| Intercept | -0.0056 | 0.017 | -0.326 | 0.745 | -0.039 | 0.028 |
| C(publication_year, Treatment(2020))[T.2000.0] | -0.0201 | 0.009 | -2.312 | 0.021 | -0.037 | -0.003 |
| C(publication_year, Treatment(2020))[T.2001.0] | -0.0262 | 0.008 | -3.206 | 0.001 | -0.042 | -0.010 |
| C(publication_year, Treatment(2020))[T.2002.0] | -0.0105 | 0.005 | -2.078 | 0.038 | -0.020 | -0.001 |
| C(publication_year, Treatment(2020))[T.2003.0] | -0.0117 | 0.004 | -3.003 | 0.003 | -0.019 | -0.004 |
| C(publication_year, Treatment(2020))[T.2004.0] | -0.0058 | 0.003 | -1.775 | 0.076 | -0.012 | 0.001 |
| C(publication_year, Treatment(2020))[T.2005.0] | -0.0017 | 0.003 | -0.532 | 0.594 | -0.008 | 0.005 |
| C(publication_year, Treatment(2020))[T.2006.0] | -0.0043 | 0.003 | -1.361 | 0.173 | -0.010 | 0.002 |
| C(publication_year, Treatment(2020))[T.2007.0] | -0.0041 | 0.003 | -1.296 | 0.195 | -0.010 | 0.002 |
| C(publication_year, Treatment(2020))[T.2008.0] | -0.0044 | 0.003 | -1.655 | 0.098 | -0.010 | 0.001 |
| C(publication_year, Treatment(2020))[T.2009.0] | -0.0112 | 0.002 | -4.486 | 0.000 | -0.016 | -0.006 |
| C(publication_year, Treatment(2020))[T.2010.0] | -0.0110 | 0.003 | -4.380 | 0.000 | -0.016 | -0.006 |
| C(publication_year, Treatment(2020))[T.2011.0] | -0.0155 | 0.002 | -6.441 | 0.000 | -0.020 | -0.011 |
| C(publication_year, Treatment(2020))[T.2012.0] | -0.0134 | 0.002 | -5.809 | 0.000 | -0.018 | -0.009 |
| C(publication_year, Treatment(2020))[T.2013.0] | -0.0220 | 0.002 | -10.165 | 0.000 | -0.026 | -0.018 |
| C(publication_year, Treatment(2020))[T.2014.0] | -0.0252 | 0.002 | -12.212 | 0.000 | -0.029 | -0.021 |
| C(publication_year, Treatment(2020))[T.2015.0] | -0.0287 | 0.002 | -14.416 | 0.000 | -0.033 | -0.025 |
| C(publication_year, Treatment(2020))[T.2016.0] | -0.0279 | 0.002 | -14.181 | 0.000 | -0.032 | -0.024 |
| C(publication_year, Treatment(2020))[T.2017.0] | -0.0262 | 0.002 | -13.631 | 0.000 | -0.030 | -0.022 |
| C(publication_year, Treatment(2020))[T.2018.0] | -0.0228 | 0.002 | -11.885 | 0.000 | -0.027 | -0.019 |
| C(publication_year, Treatment(2020))[T.2019.0] | -0.0124 | 0.002 | -6.694 | 0.000 | -0.016 | -0.009 |
| C(top_j, Treatment('OTHER'))[T.Arch.Virol] | 0.0082 | 0.006 | 1.266 | 0.206 | -0.004 | 0.021 |
| C(top_j, Treatment('OTHER'))[T.ChemRxiv] | 0.0018 | 0.005 | 0.347 | 0.729 | -0.009 | 0.012 |
| C(top_j, Treatment('OTHER'))[T.Emerg.Infect.Dis] | -0.0087 | 0.005 | -1.808 | 0.071 | -0.018 | 0.001 |
| C(top_j, Treatment('OTHER'))[T.JAMA] | -0.0297 | 0.006 | -4.730 | 0.000 | -0.042 | -0.017 |
| C(top_j, Treatment('OTHER'))[T.JMIR_Preprints] | 0.0040 | 0.006 | 0.708 | 0.479 | -0.007 | 0.015 |
| C(top_j, Treatment('OTHER'))[T.Journal_of_virology] | 0.0050 | 0.005 | 1.067 | 0.286 | -0.004 | 0.014 |
| C(top_j, Treatment('OTHER'))[T.Nature] | 0.0403 | 0.005 | 7.789 | 0.000 | 0.030 | 0.050 |
| C(top_j, Treatment('OTHER'))[T.PLoS_One] | -0.0151 | 0.003 | -4.651 | 0.000 | -0.021 | -0.009 |
| C(top_j, Treatment('OTHER'))[T.Research_Square] | 0.0031 | 0.003 | 0.983 | 0.326 | -0.003 | 0.009 |
| C(top_j, Treatment('OTHER'))[T.SSRN_Electronic_Journal] | -0.0009 | 0.003 | -0.340 | 0.734 | -0.006 | 0.004 |
| C(top_j, Treatment('OTHER'))[T.Sci_Rep] | -0.0106 | 0.005 | -2.079 | 0.038 | -0.021 | -0.001 |
| C(top_j, Treatment('OTHER'))[T.Science] | 0.0236 | 0.005 | 4.481 | 0.000 | 0.013 | 0.034 |
| C(top_j, Treatment('OTHER'))[T.Surgical_endoscopy] | -0.0055 | 0.004 | -1.249 | 0.212 | -0.014 | 0.003 |
| C(top_j, Treatment('OTHER'))[T.The_BMJ] | 0.0019 | 0.005 | 0.375 | 0.707 | -0.008 | 0.012 |
| C(top_j, Treatment('OTHER'))[T.The_Lancet] | 0.0825 | 0.006 | 13.939 | 0.000 | 0.071 | 0.094 |
| C(top_j, Treatment('OTHER'))[T.Vaccine] | -0.0183 | 0.006 | -3.204 | 0.001 | -0.029 | -0.007 |
| C(top_j, Treatment('OTHER'))[T.Virology] | -0.0051 | 0.005 | -0.945 | 0.345 | -0.016 | 0.006 |
| C(top_j, Treatment('OTHER'))[T.Viruses] | 0.0235 | 0.005 | 4.710 | 0.000 | 0.014 | 0.033 |
| C(top_j, Treatment('OTHER'))[T.bioRxiv] | -0.0053 | 0.003 | -1.602 | 0.109 | -0.012 | 0.001 |
| C(top_j, Treatment('OTHER'))[T.medRxiv] | -0.0183 | 0.002 | -8.482 | 0.000 | -0.022 | -0.014 |
| times_cited | 0.0099 | 0.000 | 25.497 | 0.000 | 0.009 | 0.011 |
| counts_mendeley | 0.0096 | 0.000 | 31.320 | 0.000 | 0.009 | 0.010 |
| counts_policy | 0.0612 | 0.002 | 29.233 | 0.000 | 0.057 | 0.065 |
| counts_twitter_unique | 0.0006 | 0.000 | 1.504 | 0.133 | -0.000 | 0.001 |
| counts_blogs_news | 0.0298 | 0.001 | 42.103 | 0.000 | 0.028 | 0.031 |
| counts_facebook | 0.0232 | 0.001 | 19.815 | 0.000 | 0.021 | 0.026 |
| expert_ratio | -0.0145 | 0.002 | -8.933 | 0.000 | -0.018 | -0.011 |
| tm_coronaviruses | -0.0055 | 0.014 | -0.399 | 0.690 | -0.032 | 0.021 |
| tm_epidemics | 0.0111 | 0.013 | 0.833 | 0.405 | -0.015 | 0.037 |
| tm_ph | -0.0025 | 0.014 | -0.183 | 0.855 | -0.029 | 0.024 |
| tm_mbi | 0.0048 | 0.013 | 0.362 | 0.717 | -0.021 | 0.031 |
| tm_clinical_medicine | -0.0145 | 0.013 | -1.089 | 0.276 | -0.041 | 0.012 |
| spectre_cluster_size | 0.0006 | 0.001 | 0.428 | 0.669 | -0.002 | 0.003 |
| network_cluster_size | -0.0005 | 0.000 | -3.816 | 0.000 | -0.001 | -0.000 |

**Table 4:** Test statistics for general topic intensities of articles cited in Wikipedia or not, limited to articles *published before 2020.* In W: cited in Wikipedia; Not in W: not cited in Wikipedia; KWH: Kruskal-Wallis H test.

| General topic | In W |  | Not in W |  | Test | p-value | Effect size |
| --- | --- | --- | --- | --- | --- | --- | --- |
|  | Mean | SD | Mean | SD | KWH | KWH | Cohen’s d |
| Coronaviruses | 0.092 | 0.156 | 0.094 | 0.165 | 0.06 | 0.807 | 0.015 |
| Epidemics | 0.225 | 0.197 | 0.161 | 0.186 | 357.84 | 0.0 | 0.341 |
| Public health | 0.195 | 0.222 | 0.172 | 0.215 | 35.158 | 0.0 | 0.107 |
| Molecular biology and immunology | 0.349 | 0.313 | 0.294 | 0.324 | 102.917 | 0.0 | 0.17 |
| Clinical medicine | 0.129 | 0.198 | 0.266 | 0.307 | 475.092 | 0.0 | 0.45 |

**Table 5:**
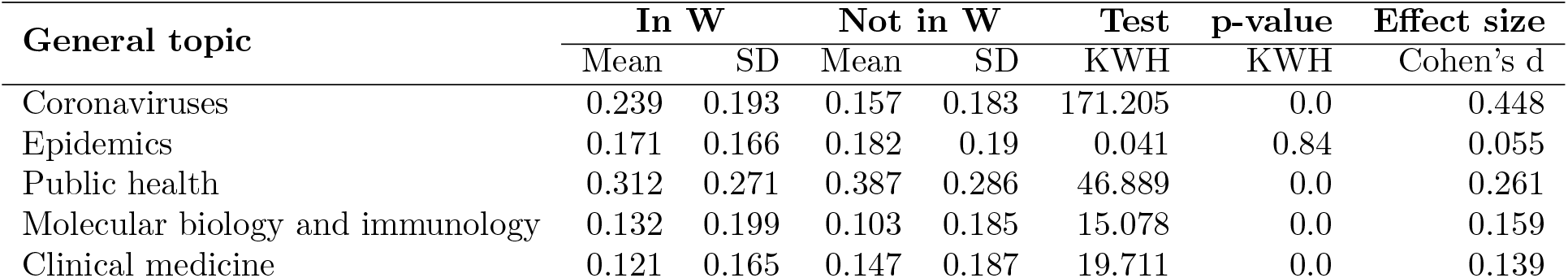
Test statistics for general topic intensities of articles cited in Wikipedia or not, limited to articles *published in 2020*. In W: cited in Wikipedia; Not in W: not cited in Wikipedia; KWH: Kruskal-Wallis H test.

**Table 6:**
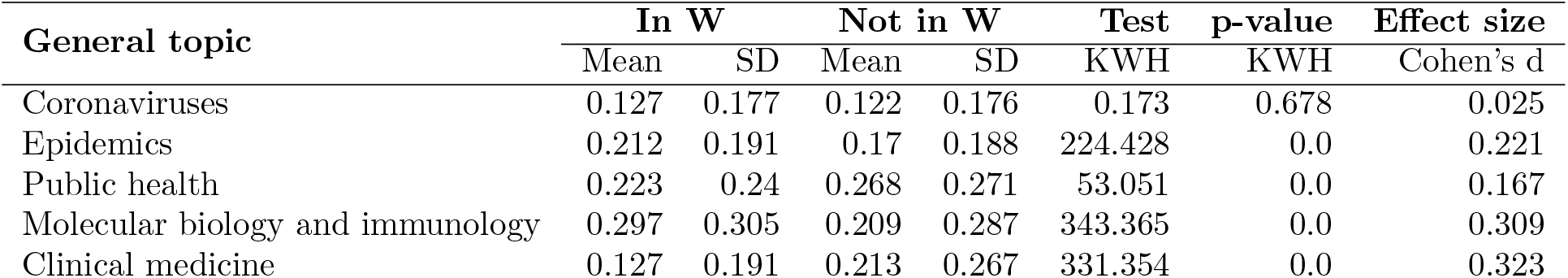
Test statistics for general topic intensities of articles cited in Wikipedia or not; *all publications*. In W: cited in Wikipedia; Not in W: not cited in Wikipedia; KWH: Kruskal-Wallis H test.

**Figure 7:**
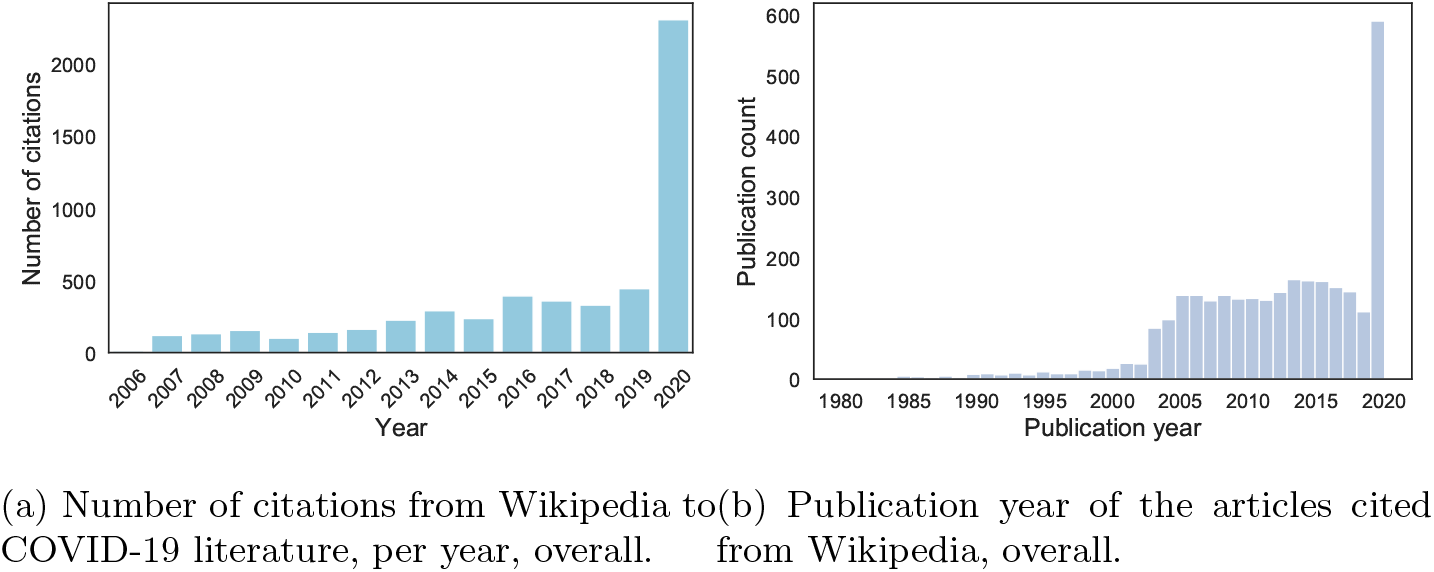
Timing of new citations from Wikipedia, and publication years of the articles they refer to.

**Figure 8:**
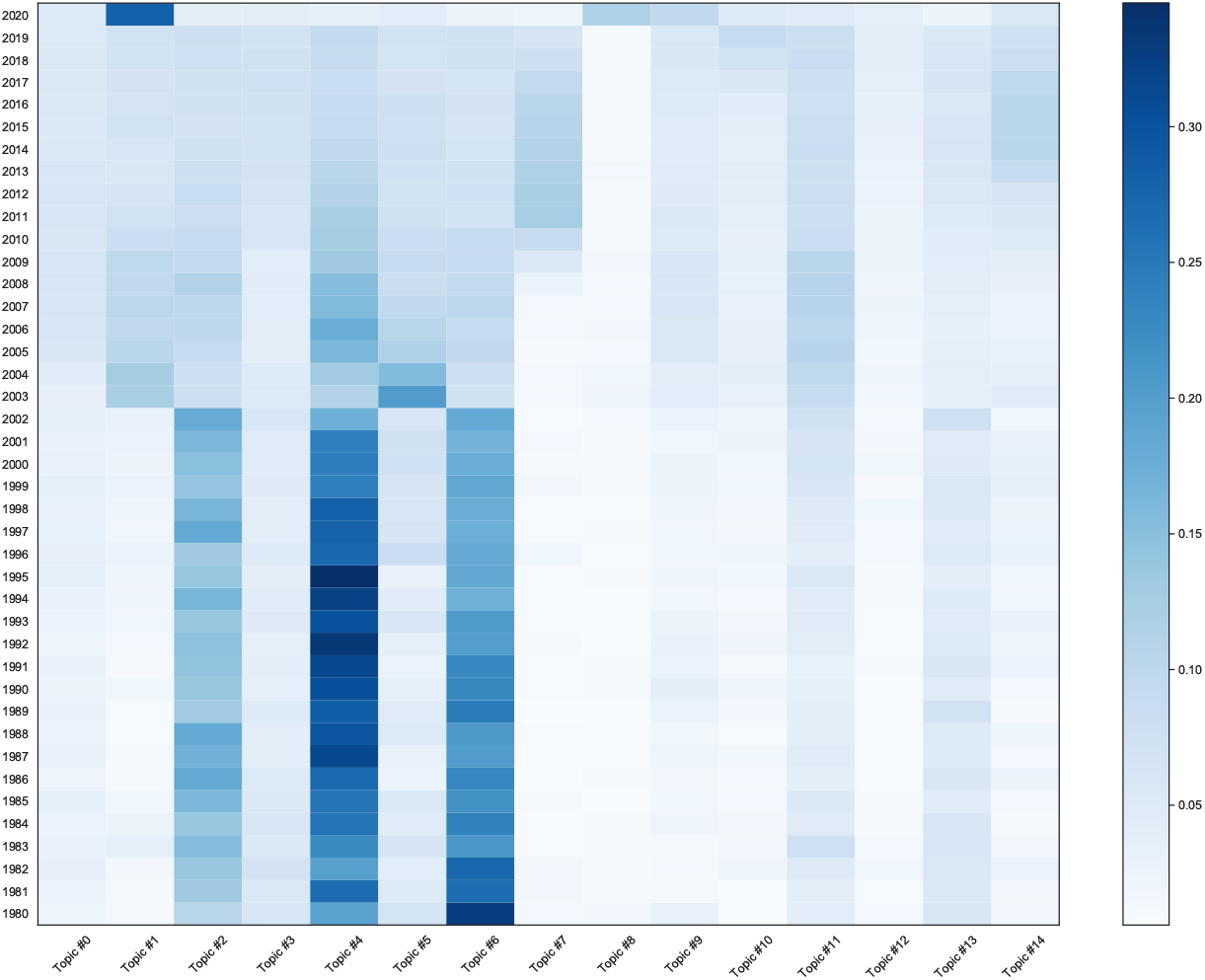
Heatmap of topic intensities over time.

**Figure 9:**
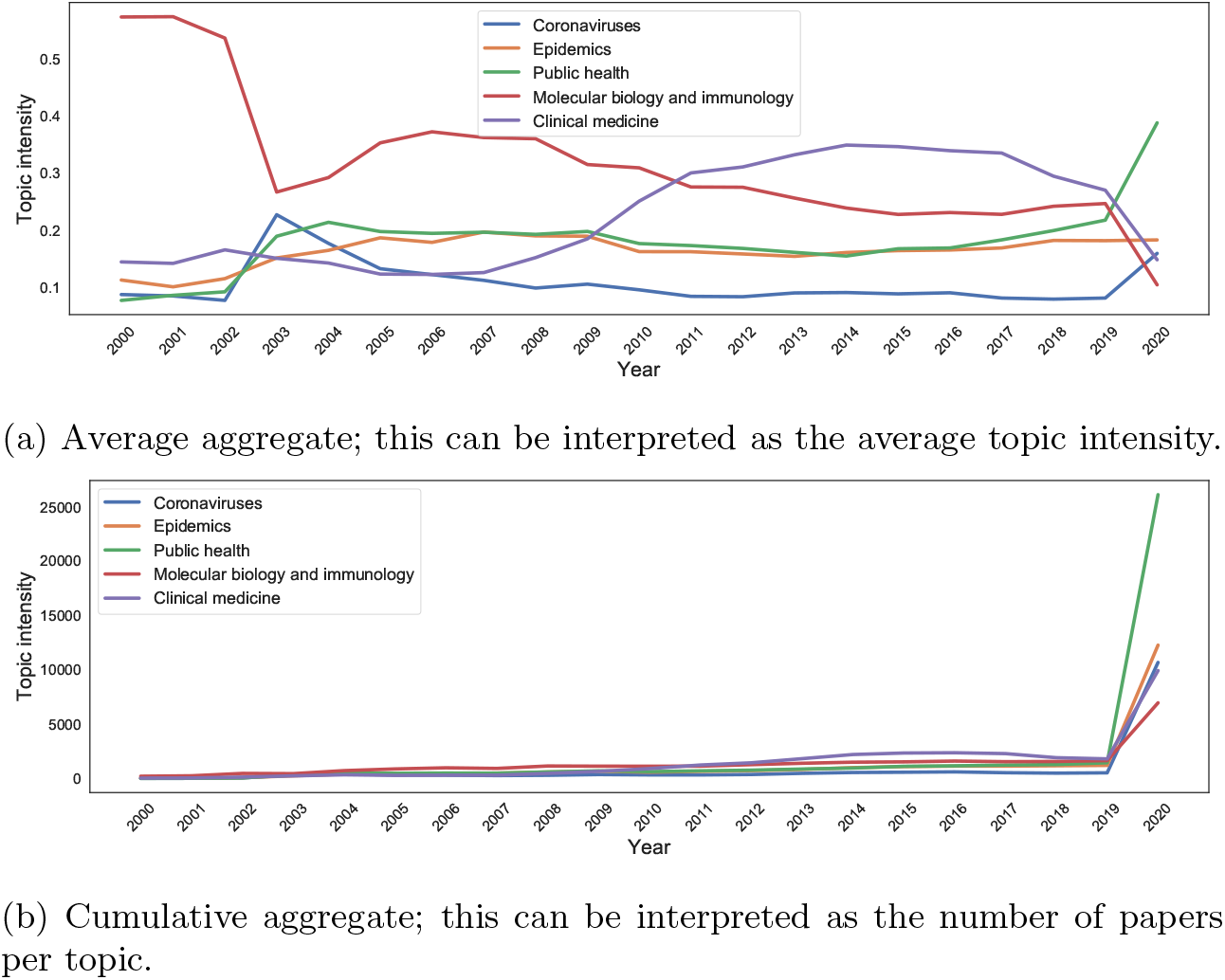
General topic intensities over time.

**Figure 10:**
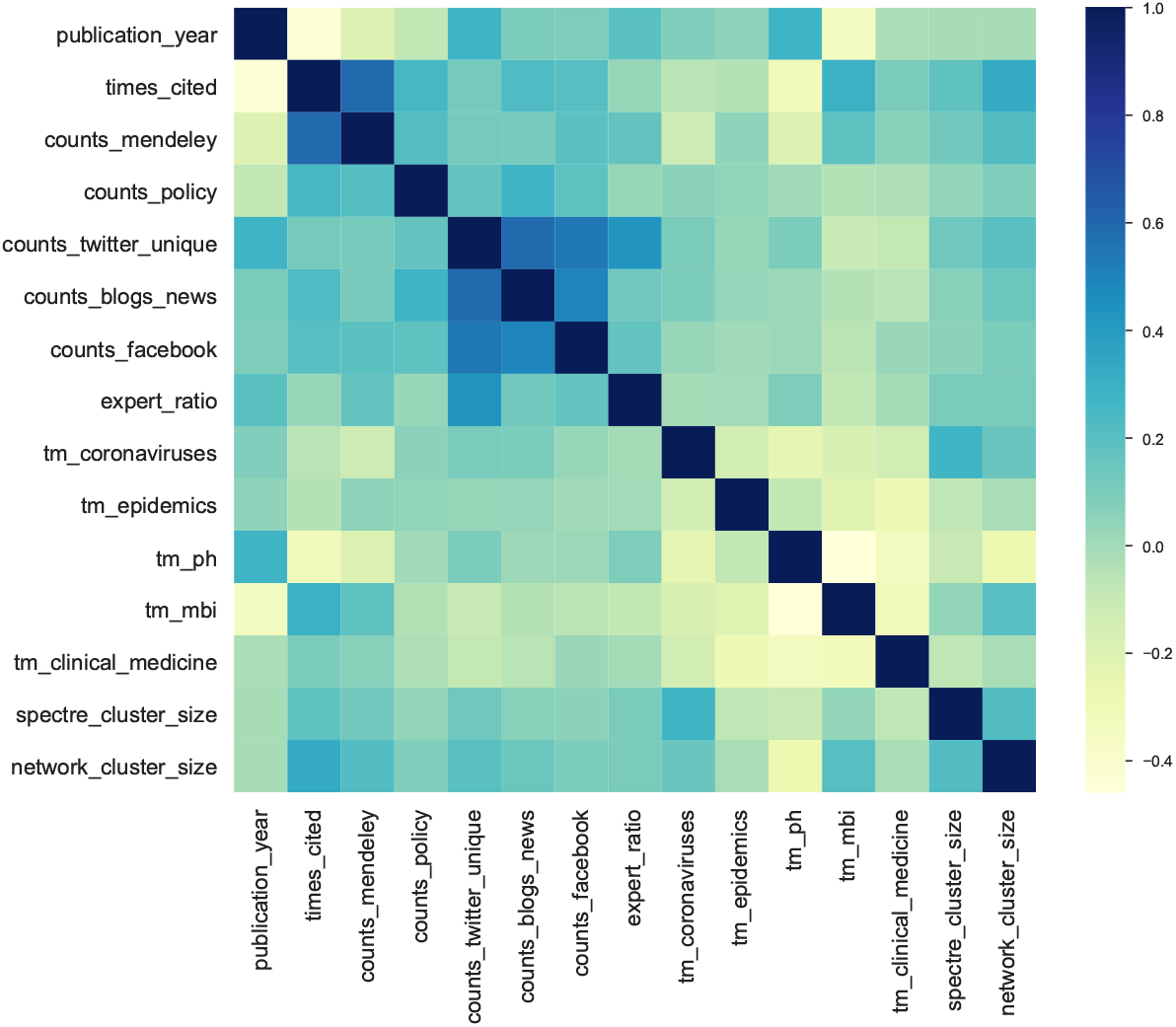
Heatmap of regression variables correlations (Pearson’s), after transformations.

**Figure 11:**
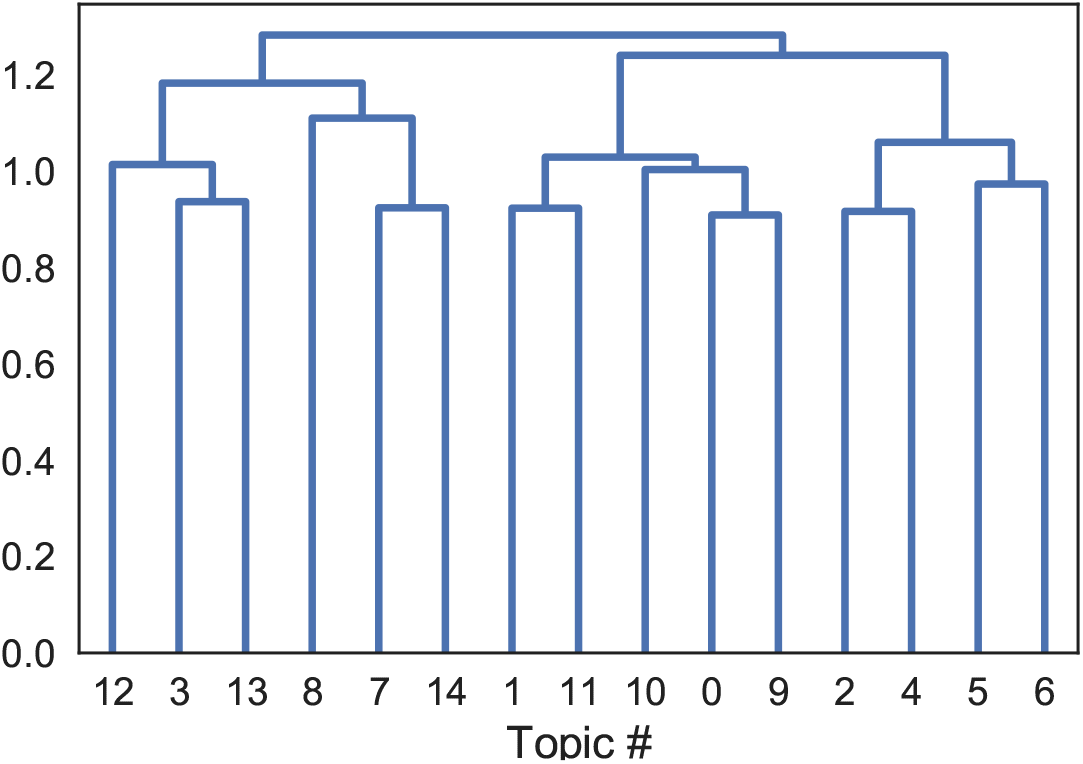
Agglomerative clustering dendrogram over topics, based on Jensen-Shannon distances. Considering a cut at 1.1, the left-most cluster (topics 3,12,13) focuses on viral epidemics and clinical medicine; next is a cluster on COVID-19 and its treatment in intensive care (topics 7,8,14); next is a cluster COVID-19, public health, epidemics and immunology (topics 0,1,9,10,11); lastly, on the right, is a cluster on molecular biology and immunology/vaccines (topics 2,4,5,6).

**Figure 12:**
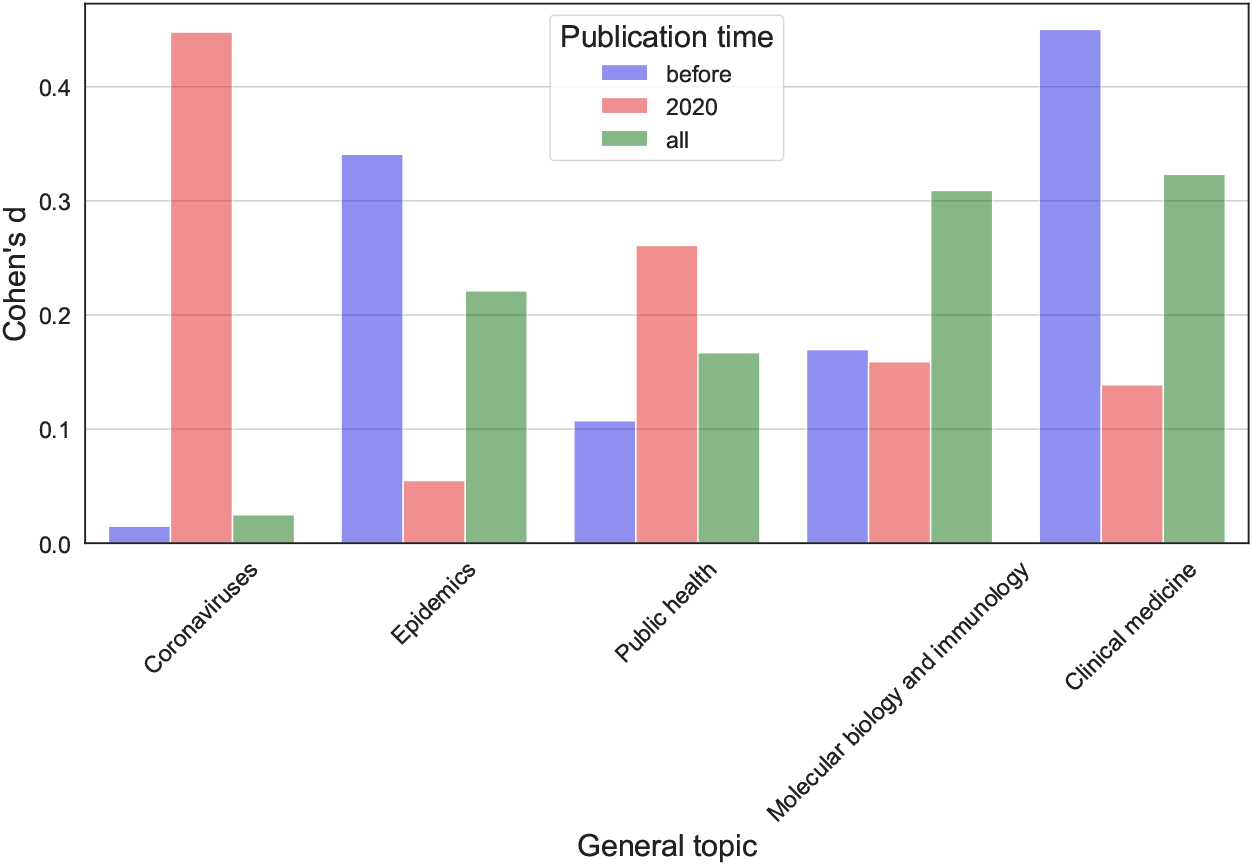
Cohen’s d effect statistic for general topic intensity differences be-tween articles cited in Wikipedia and not. Publications published before 2020, in 2020, and overall are considered. See Table 4, 5 and 6. Effect sizes are considered very small when below 0.2, small when below 0.5 and medium when below 0.8.

**Figure 13:**
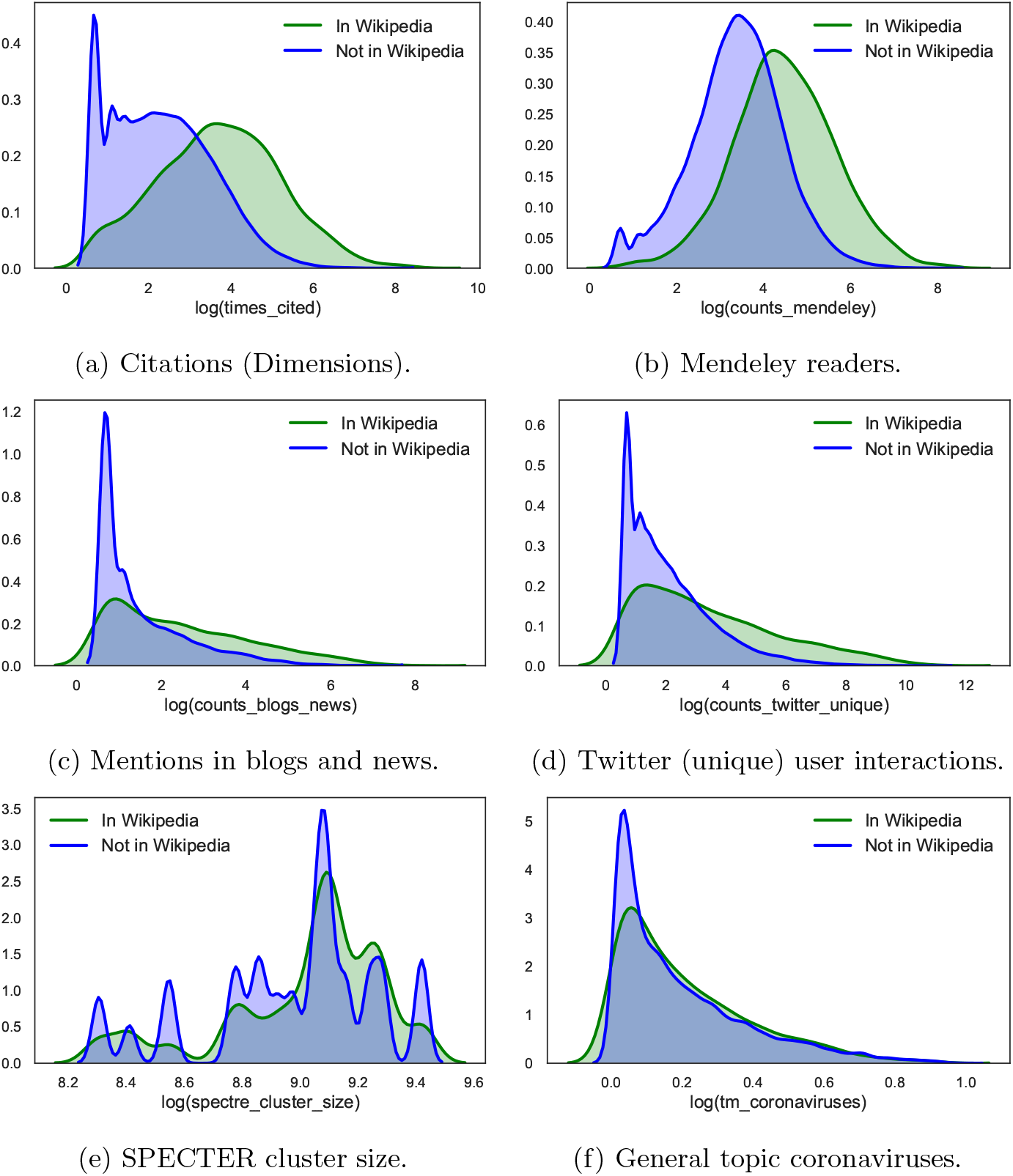
Some variables used for regression analyses. The plots distinguish variable values for articles cited from Wikipedia (green) or not (blue).

## Footnotes

1 https://wikimediafoundation.org/covid19/data [accessed 2020-07-04].

2 https://en.wikipedia.org/wiki/Wikipedia:Reliable_sources [accessed 2020-05-10].

3 https://en.wikipedia.org/wiki/Wikipedia:Identifying_reliable_sources_(medicine) [accessed 2020-05-10].

4 https://en.wikipedia.org/wiki/Wikipedia:General_sanctions [accessed 2020-05-10].

5 https://en.wikipedia.org/wiki/Wikipedia:WikiProject_COVID-19 [accessed 2020-0510].

6 The identifiers considered by Altmetric in order to establish a citation from Wikipedia to an article currently include: DOI, URI from a domain white list, PMID, PMCID, arXiv ID. https://help.altmetric.com/support/solutions/articles/6000060980-how-does-altmetric-track-mentions-on-wikipedia [accessed 2020-04-27].

7 https://github.com/allenai/paper-embedding-public-apis [accessed 2020-04-25].

8 We used gensim’s implementation for LDA [50] and tomotopy for CTM and HTM, https://bab2min.github.io/tomotopy [version 0.7.0]. The reader can find more results and the code to replicate all experiments in the accompanying repository.

9 https://en.wikipedia.org/wiki/Wikipedia:Notability [accessed 2020-05-10].

10 Calculated using Altmetric data which distinguishes among the number of researchers (*r*), experts (*e*), practitioners (*p*) and members of the public (*m*) engaging with an article. The expert ratio is defined as 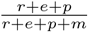.

11 Marginal effect coefficients should be interpreted as follows. For binary discrete variables (0/1), they represent the discrete rate of change in the probability of the outcome, everything else kept fix; therefore, a change from 0 to 1 with a significant coefficient of 0.01 entails an increase in the probability of the outcome of 1%. For categorical variables with more than two outcomes, they represent the difference in the predicted probabilities of any one category relative to the reference category. For continuous variables, they represent the instantaneous rate of change. It might be the case that this can also be interpreted linearly (e.g., a significant change of 1 in the variable entails a change proportional to the marginal effect coefficient in the probability of the outcome). Yet, this rests on the assumption that the relationship between independent and dependent variables is linear irrespective of the orders of magnitude under consideration. This might not be the case in practice.

## Notes

https://github.com/Giovanni1085/covid-19_wikipedia

https://github.com/CWTSLeiden/cwts_covid

### Competing Interest Statement

The authors have declared no competing interest.

### Summary of Updates

Minor revisions following a second round of reviews.

## References

[1] Dimensions COVID-19 Publications, 2020. URL: https://docs.google.com/spreadsheets/d/1-kTZJZ1GAhJ2m4GAIhw1ZdlgO46JpvX0ZQa232VWRmw/edit#gid=2034285255.

[2] EPI-WIN: WHO Information Network for Epidemics, 2020. URL: https://www.who.int/teams/risk-communication.

[3] Fighting Disinformation - Official Sources on COVID-19 - Consilium, 2020. URL: https://www.consilium.europa.eu/en/policies/covid-19-coronavirus-outbreak/fighting-disinformation.

[4] WHO COVID-19 Database, 2020. URL: https://www.who.int/emergencies/diseases/novel-coronavirus-2019/global-research-on-novel-coronavirus-2019-ncov.

[5] Clive E Adams, Alan A Montgomery, Tony Aburrow, Sophie Bloomfield, Paul M Briley, Ebun Carew, Suravi Chatterjee-Woolman, Ghalia Feddah, Johannes Friedel, Josh Gibbard, Euan Haynes, Mohsin Hussein, Mahesh Jayaram, Samuel Naylor, Luke Perry, Lena Schmidt, Umer Siddique, Ayla Serena Tabaksert, Douglas Taylor, Aarti Velani, Douglas White, and Jun Xia. Adding evidence of the effects of treatments into relevant Wikipedia pages: A randomised trial. BMJ Open, 10(2):e033655, February 2020. URL: http://bmjopen.bmj.com/lookup/doi/10.1136/bmjopen-2019-033655, doi:10.1136/bmjopen-2019-033655.

[6] Wenceslao Arroyo-Machado, Daniel Torres-Salinas, Enrique Herrera-Viedma, and Esteban Romero-Frias. Science through Wikipedia: A novel representation of open knowledge through co-citation networks. PLOS ONE, 15(2):e0228713, February 2020. URL: https://dx.plos.org/10.1371/journal.pone.0228713, doi:10.1371/journal.pone.0228713.

[7] David M. Blei. Probabilistic topic models. Communications of the ACM, 55(4):77–84, 2012. URL: http://dl.acm.org/citation.cfm?id=2133826.

[8] David M. Blei and John D. Lafferty. A correlated topic model of Science. Annals of Applied Statistics, 1(1):17–35, 2007. URL: http://projecteuclid.org/euclid.aoas/1183143727, doi:10.1214/07-AOAS114.

[9] David M. Blei, Andrew Y. Ng, and Michael I. Jordan. Latent Dirichlet allocation. Journal of Machine Learning Research, 3:993–1022, 2003. URL: http://dl.acm.org/citation.cfm?id=944937.

[10] Aleksandar Brezar and James Heilman. Readability of English Wikipedia’s health information over time. WikiJournal of Medicine, 6(1):7, 2019. URL: https://en.wikiversity.org/wiki/WikiJournal_of_Medicine/Readability_of_English_Wikipedia's_health_information_over_time, doi:10.15347/wjm/2019.007.

[11] Chih-Chun Chen and Camille Roth. {{citation needed}}: the dynamics of referencing in Wikipedia. In Proceedings of the Eighth Annual International Symposium on Wikis and Open Collaboration, Linz, Austria, 2012. ACM Press. URL: http://dl.acm.org/citation.cfm?doid=2462932.2462943, doi:10.1145/2462932.2462943.

[12] Matteo Cinelli, Walter Quattrociocchi, Alessandro Galeazzi, Carlo Michele Valensise, Emanuele Brugnoli, Ana Lucia Schmidt, Paola Zola, Fabiana Zollo, and Antonio Scala. The COVID-19 social media infodemic. arXiv:2003.05004 [nlin, physics:physics], March 2020. arXiv: 2003.05004. URL: http://arxiv.org/abs/2003.05004.

[13] Arman Cohan, Sergey Feldman, Iz Beltagy, Doug Downey, and Daniel S. Weld. SPECTER: Document-level Representation Learning using Citation-informed Transformers. arXiv:2004.07180 [cs], April 2020. arXiv: 2004.07180. URL: http://arxiv.org/abs/2004.07180.

[14] Jacob Cohen. Statistical Power Analysis for the Behavioral Sciences. Routledge, 1988. OCLC: 868313521.

[15] Giovanni Colavizza, Rodrigo Costas, Vincent A Traag, Nees Jan van Eck, Thed van Leeuwen, and Ludo Waltman. A scientometric overview of CORD-19. bioRxiv, April 2020. URL: https://www.biorxiv.org/content/10.1101/2020.04.20.046144v1, doi:10.1101/2020.04.20.046144.

[16] Zhichao Fang and Rodrigo Costas. Studying the accumulation velocity of altmetric data tracked by Altmetric.com. Scientometrics, March 2020. URL: http://link.springer.com/10.1007/s11192-020-03405-9, doi: 10.1007/s11192-020-03405-9.

[17] Andrea Forte, Nazanin Andalibi, Tim Gorichanaz, Meen Chul Kim, Thomas Park, and Aaron Halfaker. Information Fortification: An Online Citation Behavior. In Proceedings of the 2018 ACM Conference on Supporting Groupwork - GROUP ‘18, pages 83–92, Sanibel Island, Florida, USA, 2018. ACM Press. URL: http://dl.acm.org/citation.cfm?doid=3148330.3148347, doi:10.1145/3148330.3148347.

[18] Wikimedia Foundation. Responding to COVID-19. How we can help in this time of uncertainty, 2020. URL: https://wikimediafoundation.org/covid19.

[19] R. Stuart Geiger and Aaron Halfaker. When the levee breaks: without bots, what happens to Wikipedia’s quality control processes? In Proceedings of the 9th International Symposium on Open Collaboration, pages 1–6, Hong Kong, China, 2013. ACM Press. URL: http://dl.acm.org/citation.cfm?doid=2491055.2491061, doi:10.1145/2491055.2491061.

[20] Aaron Halfaker, Bahodir Mansurov, Miriam Redi, and Dario Taraborelli. Citations with identifiers in Wikipedia, 2018. URL: https://figshare.com/articles/Citations_with_identifiers_in_Wikipedia/1299540/1, doi:10.6084/m9.figshare.1299540.

[21] James M Heilman, Eckhard Kemmann, Michael Bonert, Anwesh Chatterjee, Brent Ragar, Graham M Beards, David J Iberri, Matthew Harvey, Brendan Thomas, Wouter Stomp, Michael F Martone, Daniel J Lodge, Andrea Vondracek, Jacob F de Wolff, Casimir Liber, Samir C Grover, Tim J Vickers, Bertalan Meskó, and Michael R Laurent. Wikipedia: A Key Tool for Global Public Health Promotion. Journal of Medical Internet Research, 13(1):e14, 2011. URL:http://www.jmir.org/2011/1/e14/, doi:10.2196/jmir.1589.

[22] Christian Herzog, Daniel Hook, and Stacy Konkiel. Dimensions: Bringing down barriers between scientometricians and data. Quantitative Science Studies, 1(1):387–395, February 2020. URL: https://www.mitpressjournals.org/doi/abs/10.1162/qss_a_00020, doi:10.1162/qss_a_00020.

[23] John P.A. Ioannidis. Coronavirus disease 2019: The harms of exaggerated information and non-evidence-based measures. European Journal of Clinical Investigation, page e13222, March 2020. URL: http://doi.wiley.com/10.1111/eci.13222, doi:10.1111/eci.13222.

[24] Changwook Jung, Sun Geng, Meeyoung Cha, Inho Hong, and Diego Saiez-Trumper. Open data and COVID-19: Wikipedia as an informational resource during the pandemic, 2020. URL: https://medium.com/@diegosaeztrumper/open-data-and-covid-19-wikipedia-as-an-informational-resource-during-the-pandemic-dcca

[25] Brian Keegan, Darren Gergle, and Noshir Contractor. Hot off the wiki: dynamics, practices, and structures in Wikipedia’s coverage of the Tōhoku catastrophes. In Proceedings of the 7th International Symposium on Wikis and Open Collaboration - WikiSym ‘11, Mountain View, California, 2011. ACM Press. URL: http://dl.acm.org/citation.cfm?doid=2038558.2038577, doi:10.1145/2038558.2038577.

[26] M. M. Kessler. Bibliographic coupling between scientific papers. American Documentation, 14(1):10–25, January 1963. URL: http://doi.wiley.com/10.1002/asi.5090140103, doi:10.1002/asi.5090140103.

[27] Kayvan Kousha and Mike Thelwall. Are wikipedia citations important evidence of the impact of scholarly articles and books? Journal of the Association for Information Science and Technology, 68(3):762–779, 2017. URL: http://doi.wiley.com/10.1002/asi.23694, doi:10.1002/asi.23694.

[28] Kayvan Kousha and Mike Thelwall. COVID-19 publications: Database coverage, citations, readers, tweets, news, Facebook walls, Reddit posts. a,rX?v:2004.10400 [cs], 2020. URL: https://arxiv.org/abs/2004.10400.

[29] William H. Kruskal and W. Allen Wallis. Use of Ranks in One-Criterion Variance Analysis. Journal of the American Statistical Association, 47(260):583–621, December 1952. URL: http://www.tandfonline.com/doi/abs/10.1080/01621459.1952.10483441, doi:10.1080/01621459.1952.10483441.

[30] Srijan Kumar, Robert West, and Jure Leskovec. Disinformation on the Web: Impact, Characteristics, and Detection of Wikipedia Hoaxes. In Proceedings of the 25th International Conference on World Wide Web, pages 591–602, Montr&#233al, Qu&#233bec, Canada, 2016. ACM Press. URL: http://dl.acm.org/citation.cfm?doid=2872427.2883085, doi: 10.1145/2872427.2883085.

[31] M. R. Laurent and T. J. Vickers. Seeking Health Information Online: Does Wikipedia Matter? Journal of the American Medical Informatics Association, 16(4):471–479, July 2009. URL: https://academic.oup.com/jamia/article-lookup/doi/10.1197/jamia.M3059, doi:10.1197/jamia.M3059.

[32] Florian Lemmerich, Diego Sáez-Trumper, Robert West, and Leila Zia. Why the World Reads Wikipedia: Beyond English Speakers. In Proceedings of the Twelfth ACM International Conference on Web Search and Data Mining. ACM Press, 2019. URL: http://arxiv.org/abs/1812.00474, doi:10.1145/3289600.3291021.

[33] Włodzimierz Lewoniewski, Krzysztof Wecel, and Witold Abramowicz. Analysis of References Across Wikipedia Languages. In Robertas Damaševičius and Vilma Mikašytė, editors, Information and Software Technologies, volume 756, pages 561–573. Springer International Publishing, Cham, 2017. doi:10.1007/978-3-319-67642-5_47.

[34] Loet Leydesdorff and Adina Nerghes. Co-word maps and topic modeling: A comparison using small and medium-sized corpora (N < 1,000). Journal of the Association for Information Science and Technology, 68(4):1024–1035, 2017. URL: http://doi.wiley.com/10.1002/asi.23740, doi:10.1002/asi.23740.

[35] Lauren A Maggio, Ryan M Steinberg, Tiziano Piccardi, and John M Willinsky. Reader engagement with medical content on Wikipedia. eLife, 9:e52426, March 2020. URL: https://elifesciences.org/articles/52426, doi:10.7554/eLife.52426.

[36] Lauren A Maggio, John M Willinsky, Ryan M Steinberg, Daniel Mietchen, Joseph L Wass, and Ting Dong. Wikipedia as a gateway to biomedical research: The relative distribution and use of citations in the English Wikipedia. PLOS ONE, 12(12):e0190046, 2019.

[37] Alberto Martín-Martín, Mike Thelwall, and Emilio Delgado López-Cózar. Google Scholar, Microsoft Academic, Scopus, Dimensions, Web of Science, and OpenCitations’ COCI: a multidisciplinary comparison of coverage via citations. 2020. URL: https://arxiv.org/abs/2004.14329.

[38] Mostafa Mesgari, Chitu Okoli, Mohamad Mehdi, Finn Årup Nielsen, and Arto Lanamäki. “The sum of all human knowledge”: A systematic review of scholarly research on the content of Wikipedia. Journal of the Association for Information Science and Technology, 66(2):219–245, 2015. URL: http://doi.wiley.com/10.1002/asi.23172, doi:10.1002/asi.23172.

[39] David Mimno, Hanna Wallach, Edmund Talley, Miriam Leenders, and Andrew McCallum. Optimizing semantic coherence in topic models. In Proceedings of the 2011 Conference on Empirical Methods in Natural Language Processing, pages 262–272, Edinburgh, UK, 2011. ACM.

[40] Mark Neumann, Daniel King, Iz Beltagy, and Waleed Ammar. ScispaCy: Fast and robust models for biomedical natural language processing. 2019. arXiv:arXiv:1902.07669.

[41] Finn Årup Nielsen. Scientific Citations in Wikipedia. First Monday, 12, 2007.

[42] Finn ^Å^Arup Nielsen, Daniel Mietchen, and Egon Willighagen. Scholia, Scientometrics and Wikidata. In Eva Blomqvist, Katja Hose, Heiko Paul-heim, Agnieszka Lawrynowicz, Fabio Ciravegna, and Olaf Hartig, editors, The Semantic Web: ESWC 2017 Satellite Events, volume 10577, pages 237–259. Springer International Publishing, Cham, 2017. URL: http://link.springer.com/10.1007/978-3-319-70407-4_36, doi:10.1007/978-3-319-70407-4_36.

[43] José Luis Ortega. Reliability and accuracy of altmetric providers: A comparison among Altmetric.com, PlumX and Crossref Event Data. Scientometrics, 116(3):2123–2138, September 2018. URL: http://link.springer.com/10.1007/s11192-018-2838-z, doi:10.1007/s11192-018-2838-z.

[44] Leena Paakkari and Orkan Okan. COVID-19: health literacy is an underestimated problem. The Lancet Public Health, 5(5):e249–e250, May 2020. URL: https://linkinghub.elsevier.com/retrieve/pii/S2468266720300864, doi:10.1016/S2468-2667(20)30086-4.

[45] Antonio Perianes-Rodriguez, Ludo Waltman, and Nees Jan van Eck. Constructing bibliometric networks: A comparison between full and fractional counting. Journal of Informetrics, 10(4):1178–1195, November 2016. URL: http://linkinghub.elsevier.com/retrieve/pii/S1751157716302036, doi:10.1016/j.joi.2016.10.006.

[46] Tiziano Piccardi, Miriam Redi, Giovanni Colavizza, and Robert West. Quantifying Engagement with Citations on Wikipedia. In Proceedings of The Web Conference 2020, pages 2365–2376, Taipei Taiwan, April 2020. ACM. URL: https://dl.acm.org/doi/10.1145/3366423.3380300, doi:10.1145/3366423.3380300.

[47] Alessandro Piscopo and Elena Simperl. What we talk about when we talk about Wikidata quality: a literature survey. In Proceedings of the 15th International Symposium on Open Collaboration, Skövde, Sweden, 2019. ACM Press. doi:10.1145/3306446.3340822.

[48] Reid Priedhorsky, Jilin Chen, Shyong (Tony) K. Lam, Katherine Panciera, Loren Terveen, and John Riedl. Creating, destroying, and restoring value in wikipedia. In Proceedings of the 2007 international ACM conference on Conference on supporting group work, Sanibel Island, Florida, USA, 2007. ACM Press. URL: http://portal.acm.org/citation.cfm?doid=1316624.1316663, doi:10.1145/1316624.1316663.

[49] Jason Priem, Heather A. Piwowar, and Bradley M. Hemminger. Altmetrics in the Wild: Using Social Media to Explore Scholarly Impact, 2012. URL: https://arxiv.org/html/1203.4745.

[50] Radim Řehůřek and Petr Sojka. Software framework for topic modelling with large corpora. In Proceedings of the LREC 2010 Workshop on New Challenges for NLP Frameworks, pages 45–50, Valletta, Malta, May 2010. ELRA. http://is.muni.cz/publication/884893/en.

[51] Nicolás Robinson-García, Daniel Torres-Salinas, Zohreh Zahedi, and Rodrigo Costas. New data, new possibilities: Exploring the insides of Altmetric.com. El Profesional de la Informacion, 23(4):359–366, May 2014. URL: https://recyt.fecyt.es/index.php/EPI/article/view/epi.2014.jul.03, doi:10.3145/epi.2014.jul.03.

[52] Thomas Shafee, Gwinyai Masukume, Lisa Kipersztok, Diptanshu Das, Mikael Häggström, and James Heilman. Evolution of Wikipedia’s medical content: past, present and future. Journal of Epidemiology and Community Health, pages jech-2016–208601, August 2017. URL: http://jech.bmj.com/lookup/doi/10.1136/jech-2016-208601, doi:10.1136/jech-2016-208601.

[53] Xin Shuai, Zhuoren Jiang, Xiaozhong Liu, and Johan Bollen. A comparative study of academic and Wikipedia ranking. In Proceedings of the 13th ACM/IEEE-CS joint conference on Digital libraries - JCDL ‘13, Indianapolis, Indiana, USA, 2013. ACM Press. URL: http://dl.acm.org/citation.cfm?doid=2467696.2467746, doi:10.1145/2467696.2467746.

[54] Philipp Singer, Florian Lemmerich, Robert West, Leila Zia, Ellery Wulczyn, Markus Strohmaier, and Jure Leskovec. Why We Read Wikipedia. In Proceedings of the 26th International Conference on World Wide Web, pages 1591–1600, Perth, Australia, 2017. ACM Press. URL: http://dl.acm.org/citation.cfm?doid=3038912.3052716, doi: 10.1145/3038912.3052716.

[55] Denise A. Smith. Situating Wikipedia as a health information resource in various contexts: A scoping review. PLOS ONE, 15(2):e0228786, February 2020. URL:https://dx.plos.org/10.1371/journal.pone.0228786, doi:10.1371/journal.pone.0228786.

[56] Cassidy R. Sugimoto, Sam Work, Vincent Larivière, and Stefanie Haustein. Scholarly use of social media and altmetrics: A review of the literature. Journal of the Association for Information Science and Technology, 68(9):2037–2062, 2017. URL: http://doi.wiley.com/10.1002/asi.23833, doi:10.1002/asi.23833.

[57] Briony Swire-Thompson and David Lazer. Public Health and Online Misinformation: Challenges and Recommendations. Annual Review of Public Health, 41(1):433–451, April 2020. URL: https://www.annualreviews.org/doi/10.1146/annurev-publhealth-040119-094127, doi:10.1146/annurev-publhealth-040119-094127.

[58] Yee Whye Teh, Michael I Jordan, Matthew J Beal, and David M Blei. Hierarchical Dirichlet Processes. Journal of the American Statistical Association, 101(476):1566–1581, December 2006. URL:http://www.tandfonline.com/doi/abs/10.1198/016214506000000302, doi: 10.1198/016214506000000302.

[59] Misha Teplitskiy, Grace Lu, and Eamon Duede. Amplifying the impact of open access: Wikipedia and the diffusion of science. Journal of the Association for Information Science and Technology, 68(9):2116–2127, 2017. URL: http://doi.wiley.com/10.1002/asi.23687, doi:10.1002/asi.23687.

[60] Mike Thelwall. The discretised lognormal and hooked power law distributions for complete citation data: Best options for modelling and regression. Journal of Informetrics, 10(2):336–346, 2016. doi:10.1016/j.joi.2015.12.007.

[61] Mike Thelwall and Paul Wilson. Regression for citation data: An evaluation of different methods. Journal of Informetrics, 8(4):963–971, 2014. doi: 10.1016/j.joi.2014.09.011.

[62] Daniel Torres-Salinas, Esteban Romero-Frías, and Wenceslao Arroyo-Machado. Mapping the backbone of the Humanities through the eyes of Wikipedia. Journal of Informetrics, 13(3):793–803, 2019. URL: https://linkinghub.elsevier.com/retrieve/pii/S1751157718302955, doi: 10.1016/j.joi.2019.07.002.

[63] Vincent A. Traag, Paul Van Dooren, and Yurii Nesterov. Narrow scope for resolution-limit-free community detection. Physical Review E, 84(1):016114, 2011. URL: http://journals.aps.org/pre/abstract/10.1103/PhysRevE.84.016114.

[64] Vincent A. Traag, Ludo Waltman, and Nees Jan van Eck. From Louvain to Leiden: Guaranteeing well-connected communities. Scientific Reports, 9(1):5233, December 2019. URL: http://www.nature.com/articles/s41598-019-41695-z, doi:10.1038/s41598-019-41695-z.

[65] Lucy Lu Wang, Kyle Lo, Yoganand Chandrasekhar, Russell Reas, Jiangjiang Yang, Darrin Eide, Kathryn Funk, Rodney Kinney, Ziyang Liu, William Merrill, Paul Mooney, Dewey Murdick, Devvret Rishi, Jerry Sheehan, Zhihong Shen, Brandon Stilson, Alex D. Wade, Kuansan Wang, Chris Wilhelm, Boya Xie, Douglas Raymond, Daniel S. Weld, Oren Etzioni, and Sebastian Kohlmeier. CORD-19: The Covid-19 Open Research Dataset. arXiv:200f.10706 [cs], April 2020. arXiv: 2004.10706. URL: http://arxiv.org/abs/2004.10706.

[66] Bo Xie, Daqing He, Tim Mercer, Youfa Wang, Dan Wu, Kenneth R. Fleischmann, Yan Zhang, Linda H. Yoder, Keri K. Stephens, Michael Mackert, and Min K. Lee. Global health crises are also information crises: A call to action. Journal of the Association for Information Science and Technology, March 2020. URL: https://onlinelibrary.wiley.com/doi/abs/10.1002/asi.24357, doi:10.1002/asi.24357.

[67] Chyi-Kwei Yau, Alan Porter, Nils Newman, and Arho Suominen. Clustering scientific documents with topic modeling. Scientometrics, 100(3):767–786, 2014. URL: http://link.springer.com/10.1007/s11192-014-1321-8, doi:10.1007/s11192-014-1321-8.

[68] Zohreh Zahedi, Rodrigo Costas, and Paul Wouters. How well developed are altmetrics? A cross-disciplinary analysis of the presence of ‘alternative metrics’ in scientific publications. Scientometrics, 101(2):1491–1513, 2014. URL: http://link.springer.com/10.1007/s11192-014-1264-0, doi:10.1007/s11192-014-1264-0.

[69] John Zarocostas. How to fight an infodemic. Lancet, 395(10225), February 2020. URL: https://linkinghub.elsevier.com/retrieve/pii/S014067362030461X, doi:10.1016/S0140-6736(20)30461-X.

